# Collaboration between IL-7 and IL-15 enables adaptation of tissue-resident and circulating memory CD8^+^ T cells

**DOI:** 10.1101/2024.05.31.596695

**Authors:** Nicholas N. Jarjour, Talia S. Dalzell, Nicholas J. Maurice, Kelsey M. Wanhainen, Changwei Peng, Taylor A. DePauw, Katharine E. Block, William J. Valente, K. Maude Ashby, David Masopust, Stephen C. Jameson

## Abstract

Interleukin-7 (IL-7) is considered a critical regulator of memory CD8^+^ T cell homeostasis, but this is primarily based on analysis of circulating and not tissue-resident memory (T_RM_) subsets. Furthermore, the cell-intrinsic requirement for IL-7 signaling during memory homeostasis has not been directly tested. Using inducible deletion, we found that *Il7ra* loss had only a modest effect on persistence of circulating memory and T_RM_ subsets and that IL-7Rα was primarily required for normal basal proliferation. Loss of IL-15 signaling imposed heightened IL-7Rα dependence on memory CD8^+^ T cells, including T_RM_ populations previously described as IL-15-independent. In the absence of IL-15 signaling, IL-7Rα was upregulated, and loss of IL-7Rα signaling reduced proliferation in response to IL-15, suggesting cross-regulation in memory CD8^+^ T cells. Thus, across subsets and tissues, IL-7 and IL-15 act in concert to support memory CD8^+^ T cells, conferring resilience to altered availability of either cytokine.

**Highlights:** Tissue-resident and circulating memory CD8^+^ T cells modestly decline after loss of IL-7Rα IL-7Rα is required for normal self-renewal of memory CD8^+^ T cells

Combined loss of IL-7 and IL-15 causes a profound defect across memory CD8^+^ T cell subsets Cross-regulation of IL-7 and IL-15 signaling occurs in memory CD8^+^ T cells

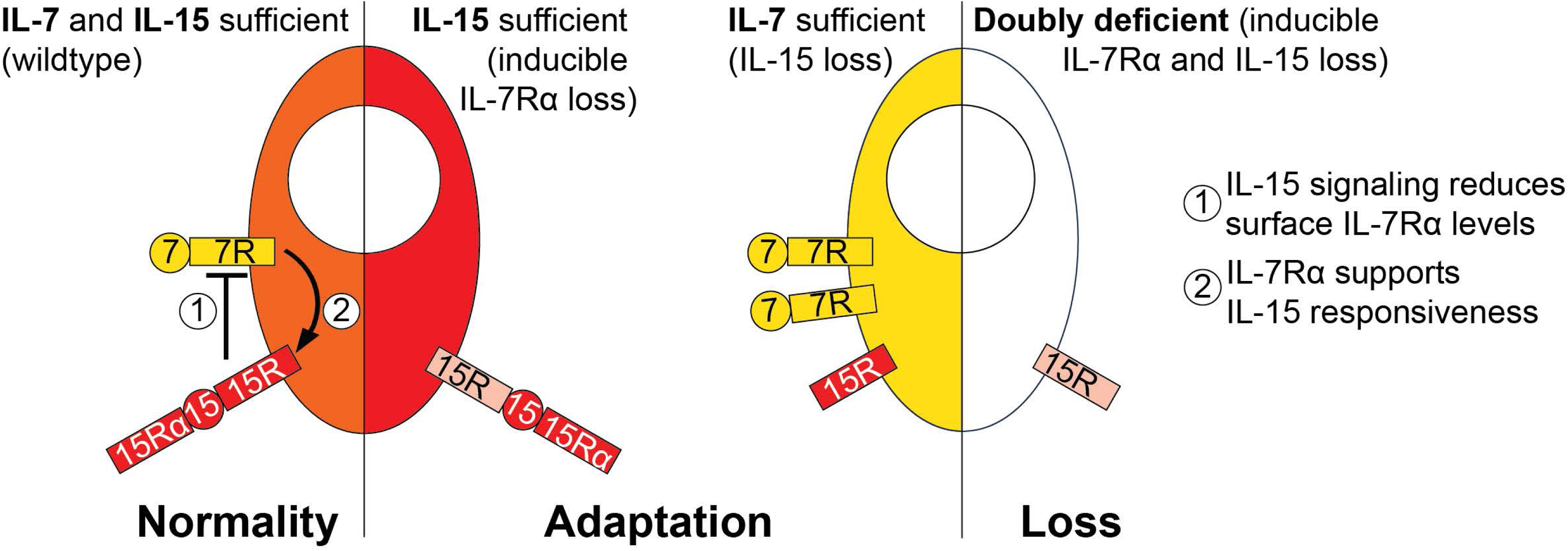

## INTRODUCTION

Classical immunological memory is formed in the wake of a successful immune response and establishes stable cellular and humoral immunity against the eliciting pathogen. Together these form the basis for a potent and protective recall response upon re-exposure to the pathogen, mediating long-term defense of the organism. CD8^+^ T cell memory was first characterized in circulating populations found in the blood and lymphoid organs, but more recently abundant tissue-resident memory CD8^+^ T cells (T_RM_) have been identified in diverse organs^1–3^. These populations are locally maintained independent of contribution from the circulation^4,5^, acquire tissue-specific gene expression programs^6–12^, and are capable of local expansion and control of infection and cancer at the point of origin^13–17^. Perhaps in part because T_RM_ adapt to their tissue environments, their requirements for generation and maintenance are known to differ from those of circulating memory populations^2,3^.

The common gamma chain-(γC)-dependent cytokines IL-7 and IL-15 are each thought to be critical for maintenance of CD8^+^ T cell memory, based largely on early studies of circulating populations^18–25^. With regard to T_RM_ requirements for IL-7 and IL-15 signaling, levels of the receptor components IL-2Rβ and IL-7Rα are known to be reduced on some T_RM_^26,27^, which could indicate lessened dependence on tonic levels of these cytokines. Indeed, more recent work has revealed that requirements for IL-15 are in fact variable for memory CD8^+^ T cells (particularly for some T_RM_), depending on the memory subset, route of infection, and number of previous recall responses^28,29^. However, we recently showed that IL-15 responsiveness is conserved across circulating and resident memory CD8^+^ T cell subsets, including populations that do not depend on IL-15 for normal homeostasis, indicating that such memory populations retain the capacity to respond to IL-15^27^. IL-7 is still considered a crucial regulator of CD8^+^ T cell memory^30–33^, though few studies have assessed this for T_RM_^34^. Notably, seminal early work used transfer of T cells derived from germline *Il7ra*^-/-^ mice and antibody blockade of IL-7 signaling to assess requirements for memory CD8^+^ T cells in the blood and lymphoid tissues^20,22^. As noted by the authors as a concern, development of T cells in *Il7ra*^-/-^ mice is severely compromised and IL-7Rα blockade may elicit lymphopenia. Global and lifelong deficiency in IL-7Rα could result in confounding defects in naïve CD8^+^ T cells that exaggerate the requirement for IL-7Rα in memory CD8^+^ T cells or, alternatively, adaptation to become IL-7Rα-independent, underestimating the true severity of IL-7Rα deficiency^20,35^. Other studies have assessed IL-7 dependence in a variety of ways, with mild to severe defects in the absence of IL-7 signaling during and after memory differentiation^35–41^. In light of these concerns, Carrette and Surh^31^ highlighted the importance of cell-intrinsically ablating IL-7 signaling during T cell memory homeostasis, without globally inhibiting IL-7 signaling. Intriguingly, IL-7Rα is regulated by IL-7 and IL-15 signaling in primarily naïve T cell populations, suggesting that loss of one signaling modality could alter the other^42^. Taken together, how IL-7 and IL-15 regulate memory CD8^+^ T cells (and especially the T_RM_ compartment) is in need of reassessment. Therefore, we set out to specifically address the role for IL-7Rα in the homeostasis of circulating and tissue-resident memory CD8^+^ T cells and to attempt to resolve how IL-7 and IL-15 jointly regulate CD8^+^ T cell memory.

Unexpectedly, inducible deletion of *Il7ra* caused relatively modest impairment of memory CD8^+^ T cell maintenance for both circulating and resident memory subsets, including T_RM_ populations that were previously described as IL-15-independent. IL-7Rα was required for normal basal proliferation of memory cells during homeostasis in most locales, but not in response to recall stimulation. Deficiency in IL-15 led to IL-7Rα upregulation and greatly augmented IL-7Rα- dependence, suggesting that IL-15-independent CD8^+^ T cell memory is explained by compensatory IL-7 signaling. Loss of signaling for both cytokines caused a pronounced defect in memory cells in all sites tested. Furthermore, loss of IL-7Rα during memory homeostasis resulted in reduced proliferation in response to IL-15 treatment, suggesting cooperativity between IL-7 and IL-15. We propose a new model for cytokine regulation of CD8^+^ T cell memory, primarily relying on integrated regulation by IL-7 and IL-15 with capacity for adaptation in the absence of either cytokine.

## RESULTS

### Inducible deletion of *Il7ra* in pre-formed memory CD8^+^ T cells has modest effects on maintenance, but alters proliferation

To assess whether IL-7Rα was stringently required for established CD8^+^ T cell memory cells, *Il7ra*^fl/fl^ mice were backcrossed and intercrossed with the tamoxifen-inducible *Ubc-cre*^ERT2^ and P14 TCR transgenic lines. Congenically distinct control (*Ubc-cre*^ERT2^ *Il7ra*^+/+^ or Cre-negative *Il7ra*^fl/fl^) and *Ubc-cre*^ERT2^ *Il7ra*^fl/fl^ P14 T cells were adoptively co-transferred to recipients, followed by infection with LCMV Armstrong to elicit differentiation into memory cells. Tamoxifen could then be given at a memory timepoint to ablate *Il7ra* in pre-formed memory P14 cells (**Figure 1A**) and specifically interrogate the importance of IL-7Rα during memory homeostasis. Co-transfer was used so that (after administration of tamoxifen) IL-7Rα-sufficient and -deficient P14 T cells could compete within the same recipient, increasing sensitivity to detect an advantage for IL-7Rα- sufficient memory cells. First, deletion was confirmed via staining for CD127 (IL-7Rα), revealing robust tamoxifen-inducible loss of CD127 on *Ubc-cre*^ERT2^ *Il7ra*^fl/fl^ P14 T cells across tissues, as well as generally lower CD127 expression by control P14 T cells in non-lymphoid tissues (most clearly for the IEL, SG, and FRT) (**Figures 1B-1D**)^26^. Loss of CD127 was essentially completed within a week of starting tamoxifen treatment (data not shown). Nevertheless, while we expected a profound defect to appear in the absence of IL-7Rα, at least for circulating memory populations^20^, loss of *Il7ra* only conferred a minor competitive disadvantage (∼2-3 fold by ∼2 months post-tamoxifen) and did not result in selective loss of lymphoid or nonlymphoid tissue (NLT) memory populations (**Figures 1E, S1A, and S1B**). This is a notable contrast with IL-15 deficiency, which has been reported to cause substantial defects in memory CD8^+^ T cell homeostasis across lymphoid tissues and several non-lymphoid tissue sites, although loss of IL- 15 has very little effect on T_RM_ in some sites (such as small intestine and female reproductive tract)^28^. One possible explanation for IL-15-independent memory CD8^+^ T cell populations is that IL-15-dependent sites were largely IL-7-independent, and IL-15-independent sites (IEL, FRT) were highly IL-7-dependent. However, this simplistic explanation did not hold true in our hands, indicating that cytokine regulation of CD8^+^ T cell memory has an additional layer of complexity.

**Figure 1:**
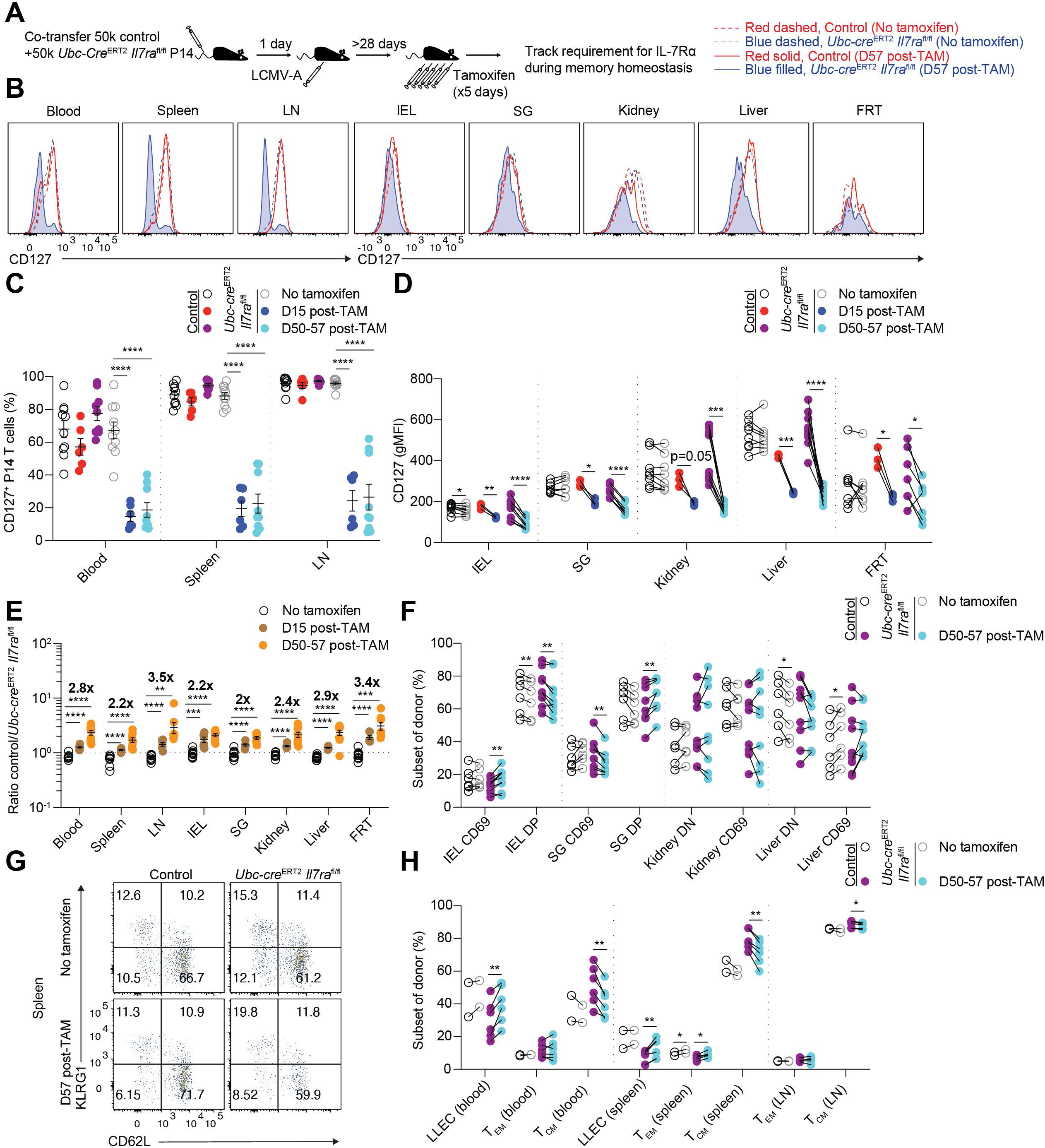
Circulating and resident memory P14 T cells are resilient to loss of *Il7ra* during memory homeostasis. (A) Congenic control (*Ubc-cre*^ERT2^ *Il7ra*^+/+^ or *Il7ra*^fl/fl^) and *Ubc-cre*^ERT2^ *Il7ra*^fl/fl^ P14 T cells were co-transferred to recipients, followed by LCMV-Armstrong infection one day later. After resting to memory phase (>28 days), some mice were given i.p. tamoxifen for 5 consecutive days to initiate Cre-mediated recombination of the floxed *Il7ra* allele and tracked over time via congenic markers. (B) Representative flow cytometry for CD127 (IL-7Rα) on control and *Ubc-cre*^ERT2^ *Il7ra*^fl/fl^ memory P14 T cells and quantitation for (C) percent CD127^+^ and (D) CD127 gMFI as in (B) from untreated and tamoxifen-treated mice. (E) Quantitation of the ratio of control and *Ubc-cre*^ERT2^ *Il7ra*^fl/fl^ P14 T cells from untreated and tamoxifen-treated mice. (F) Quantitation of the proportions of NLT memory P14 subsets (DN, CD69^-^ CD103^-^. DP CD69^+^ CD103^+^) from each donor from untreated and tamoxifen-treated mice. (G) Representative flow cytometry for CD62L and KLRG1 for splenic memory P14 T cells and (H) quantitation of the proportion of long-lived effector cells (LLEC, KLRG1^+^ CD62L^-^), T effector memory (T_EM_ KLRG1^-^ CD62L^-^), and T central memory (T_CM_ KLRG1^-^CD62L^+^) from each donor from untreated and tamoxifen-treated mice as in (G). LN, inguinal LN. IEL, small intestine intraepithelial lymphocytes. SG, salivary gland. FRT, female reproductive tract. Data are (B,G) representative of 3 experiments (n=6-9/group), (C,E,F,H) compiled from 2-5 experiments (n=4-10/group), or (D) compiled from 1-5 experiments (n=3-10/group). Error is expressed as ± S.E.M. Unpaired (C,E) and paired Student’s t tests (D,F,H).* p<0.05. ** p<0.01. *** p<0.001. **** p<0.0001. See also Figures S1 and S2.

To compare IL-7Rα dependence of established memory CD8^+^ T cells to that of a population known to be highly IL-7Rα-dependent^20,43,44^, naïve control and *Ubc-cre*^ERT2^ *Il7ra*^fl/fl^ P14 T cells were co-transferred into naïve recipients (**Figure S2A**). In tamoxifen-treated mice, induced *Il7ra*-deficient naïve P14 T cells were rapidly lost and exhibited a profound competitive disadvantage (>25 fold by ∼2 weeks post-tamoxifen), as previously described^44^ and in stark contrast with memory P14 T cells (**Figures 1E versus S2B and S2C**). By two weeks after tamoxifen treatment, remaining *Ubc-cre*^ERT2^ *Il7ra*^fl/fl^ naïve CD8^+^ T cells were largely CD127- positive escapees of Cre-mediated recombination, in contrast to memory P14 T cells, which largely remained CD127-negative (**Figures 1C and 1D versus S2D**). Therefore, memory CD8^+^ T cells exhibited a modest dependence on IL-7Rα across subsets and far greater resilience than naïve CD8^+^ T cells to loss of IL-7Rα.

After administration of tamoxifen, *Il7ra*-deficient P14 T cells showed relatively normal proportions of T_RM_ phenotype subsets, but a slightly higher proportion of KLRG1^+^ long-lived effector cells (LLEC) and a slightly lower proportion of CD62L^+^ central memory cells (T_CM_) (**Figures 1F-1H, S1C, and S1D**). In all experiments, intravenous labelling was used to discriminate vascular and tissue-localized memory cells^45^, while CD69/CD103 expression was also used to identify T_RM_^4,27^. No clear differences were observed when assessing necrotic/apoptotic cells via Annexin V (**Figure S1E**). However, consistent with previous reports^20,39,40^, Bcl2 staining was reduced for circulating memory cells in the absence of IL-7Rα, though Bcl2 levels of IL-7Rα-deficient memory cells remained higher than those of naïve T cells (**Figures S1F and S1G**)^46^. Taken together, induced ablation of *Il7ra* in pre-formed memory CD8^+^ T cells did not result in a strong predisposition towards cell death.

However, when proliferation of memory P14 T cells was assessed after ablation of *Il7ra*, we observed a consistent reduction in the proportion of cycling cells across circulating memory populations using both Ki67 staining and BrdU incorporation (as an indicator of DNA synthesis during S phase) (**Figures 2A-2D, S3A, and S3B)**. Proliferation was also reduced for most NLT populations, but memory P14 T cells in the SG and IEL were only modestly affected (**Fig. 2A-2D, S3A, and S3B**). For proliferation of specific memory subsets, induced loss of *Il7ra* affected all circulating memory populations, but again had a more complex effect on NLT-localized populations (**Figures S3C-F**). However, generally consistent trends were observed for resident phenotype (CD69-positive ± CD103-positive) and circulating phenotype memory P14 T cells within the same tissue. Taken together, our data indicate that IL-7Rα regulates self-renewal of memory CD8^+^ T cells, with the most pronounced effect on circulating memory populations.

**Figure 2:**
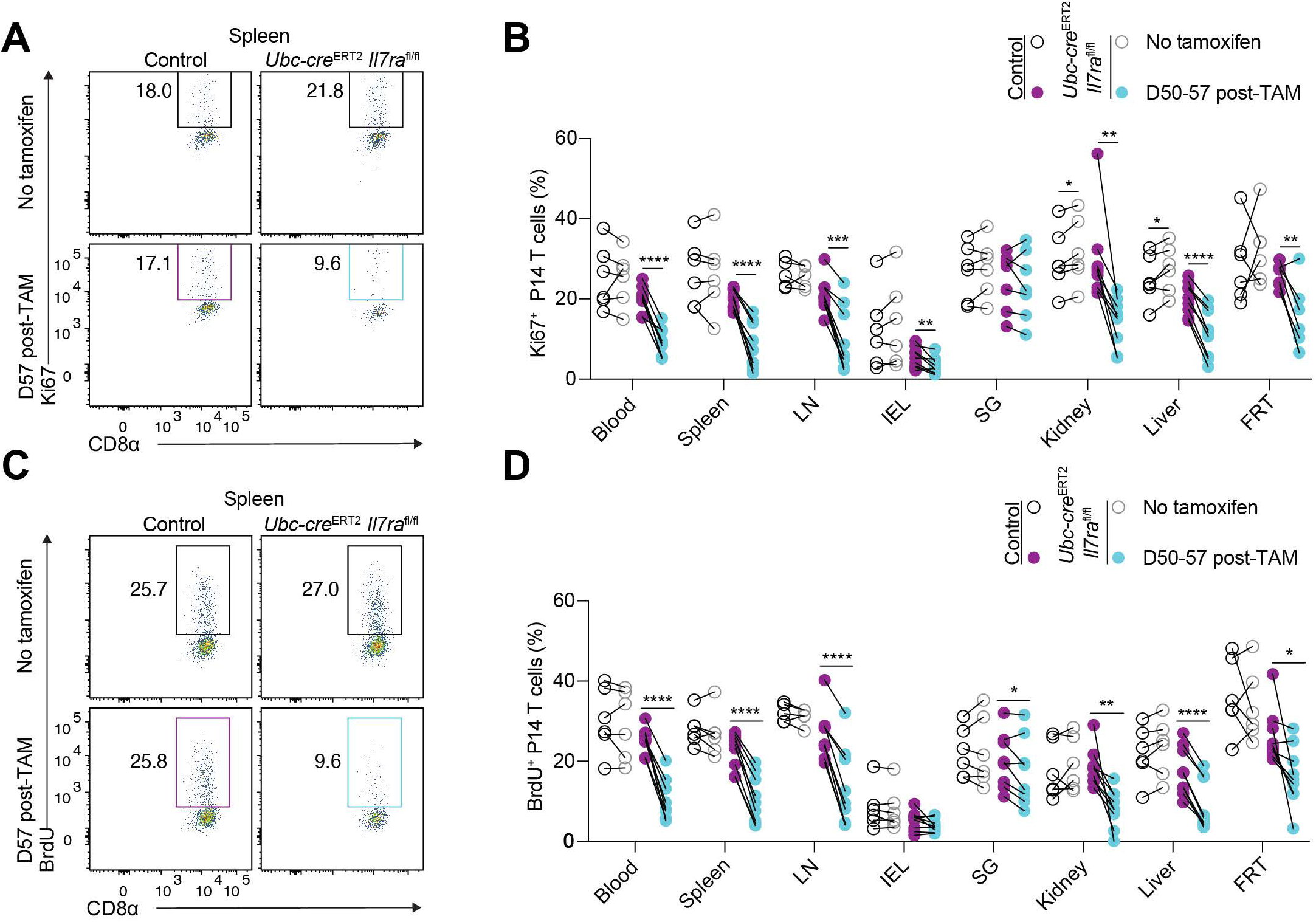
Loss of *Il7ra* alters proliferation of memory CD8^+^ T cell subsets. Congenic control (*Ubc-cre*^ERT2^ *Il7ra*^+/+^ or *Il7ra*^fl/fl^) and *Ubc-cre*^ERT2^ *Il7ra*^fl/fl^ P14 T cells were co-transferred to recipients, followed by LCMV-Armstrong infection one day later. After resting to memory phase (>28 days), some mice were given i.p. tamoxifen for 5 consecutive days to initiate Cre-mediated recombination of the floxed *Il7ra* allele. (A) Representative flow cytometry for Ki67 expression and (B) quantitation of the proportion of Ki67-expressing control and *Ubc-cre*^ERT2^ *Il7ra*^fl/fl^ memory P14 T cells as in (A) from untreated and tamoxifen-treated mice. (C) Representative flow cytometry for BrdU incorporation and (D) quantitation of the proportion of BrdU-incorporating control and *Ubc-cre*^ERT2^ *Il7ra*^fl/fl^ memory P14 T cells as in (C) from untreated and tamoxifen-treated mice. Data are (A,C) representative of 3 experiments (n=5-9/group) or (B,D) compiled from 3-5 experiments (n=5-9/group). Error is expressed as ± S.E.M. Paired Student’s t tests. * p<0.05. ** p<0.01. *** p<0.001. **** p<0.0001. See also Figures S1 and S3.

As P14 TCR transgenic T cells could be unusual in their IL-7-dependence, we also assessed the impact of induced *Il7ra* loss from LCMV-specific polyclonal CD8^+^ T cells. Mixed bone marrow chimeras were generated using congenically distinct control (*Il7ra*^fl/fl^) and *Ubc- cre*^ERT2^ *Il7ra*^fl/fl^ marrow transferred into irradiated wildtype recipients. After reconstitution, these animals were infected with LCMV to elicit antigen-specific memory populations trackable with peptide/MHC tetramers for the D^b^-restricted gp33, gp276, and NP396 epitopes of LCMV (**Figure 3A**). After establishment of memory populations, tamoxifen was given to delete *Il7ra*, allowing us to track the effects of IL-7Rα deficiency for these oligoclonal populations of different specificities. Employing mixed chimeras allowed us to assess direct competition between IL-7Rα-sufficient and -deficient naïve and memory CD8^+^ T cells as well as to mitigate the impact of potential lymphopenia (due to loss of IL-7Rα) via control bone marrow cells.

**Figure 3:**
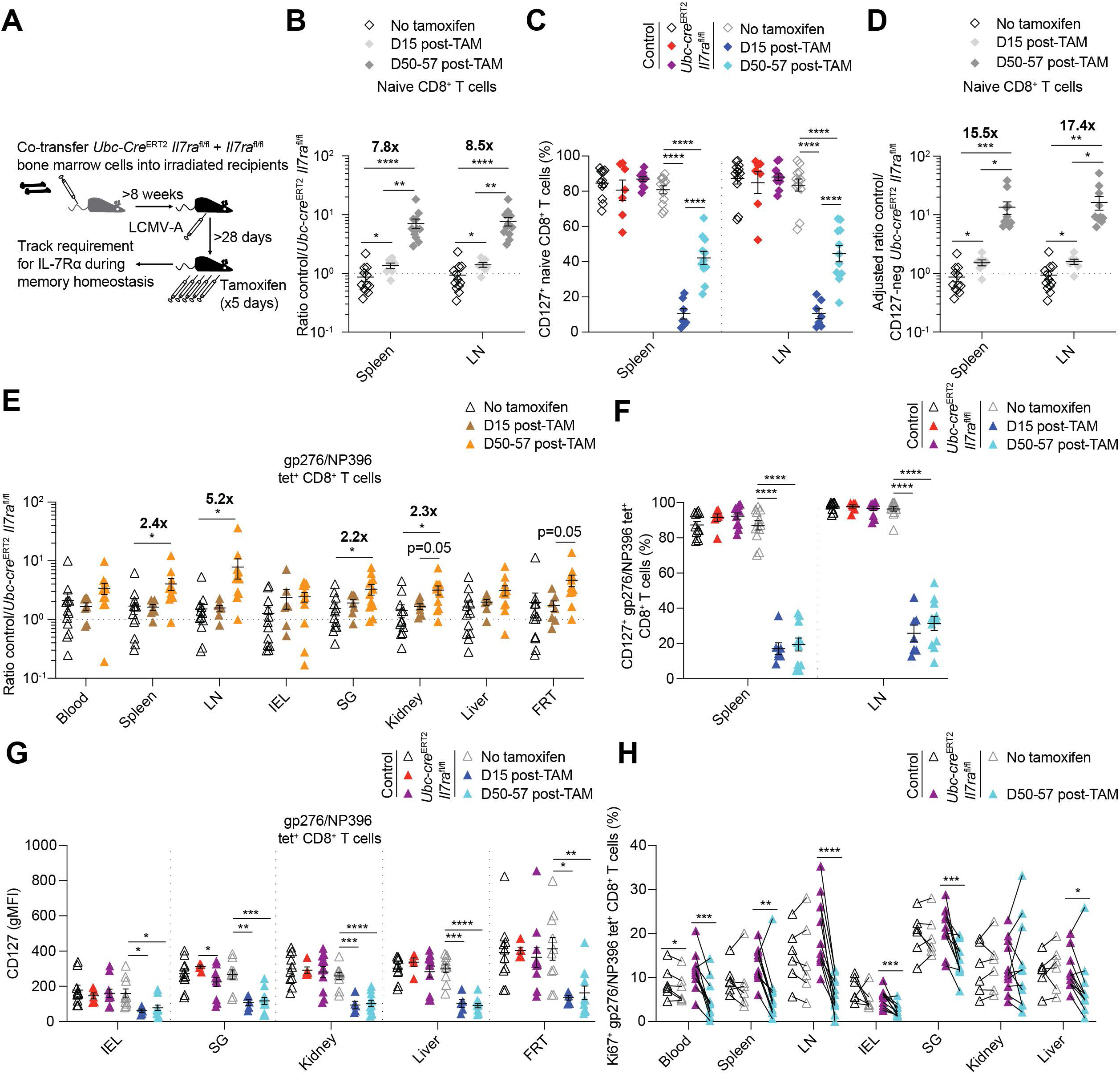
Oligoclonal memory CD8^+^ T cells of defined antigen specificity are resilient to loss of *Il7ra* during memory homeostasis. (A) Congenic *Il7ra*^fl/fl^ *Ubc-cre*^ERT2^-negative (control) and -positive bone marrow was co-transferred to lethally irradiated recipients. After >8 weeks of reconstitution, animals were infected with LCMV- Armstrong. After resting to memory phase (>28 days), some mice were given i.p. tamoxifen for 5 consecutive days to initiate Cre-mediated recombination of the floxed *Il7ra* allele and tracked over time via tetramer staining and congenic markers. (B-D) Quantitation of control and *Ubc-cre*^ERT2^ *Il7ra*^fl/fl^ naïve (CD44^lo^ CD62L^+^) CD8^+^ T cells for (B) ratio, (C) proportion CD127^+^, and (D) ratio of control to CD127^-^ *Ubc-cre*^ERT2^ *Il7ra*^fl/fl^ naïve cells from untreated and tamoxifen-treated bone marrow chimeras. (E) Quantitation of the ratio of control and *Ubc-cre*^ERT2^ *Il7ra*^fl/fl^ gp276/D^b^/NP396/D^b^ tetramer-binding memory cells from untreated and tamoxifen-treated bone marrow chimeras. (F-H) Quantitation of the (F) proportion CD127^+^, (G) CD127 gMFI, and (H) proportion of Ki67-expressing cells for control and *Ubc-cre*^ERT2^ *Il7ra*^fl/fl^ gp276/D^b^/NP396/D^b^ tetramer-binding memory cells from untreated and tamoxifen-treated bone marrow chimeras. Data are compiled from 3-7 experiments (n=6-12/group). Error is expressed as ± S.E.M. Unpaired (B-G) and paired Student’s t tests (H). * p<0.05. ** p<0.01. *** p<0.001. **** p<0.0001. See also Figure S4.

As expected^20,43,44^ and as we observed in the P14 system, loss of *Il7ra* conferred a disadvantage to naïve CD44^lo^ CD62L^+^ polyclonal CD8^+^ T cells, which also experienced strong selection for IL-7Rα (CD127)-expressing escapees of *Il7ra* deletion (**Figures 3B and 3C**). When the ratio of control *Il7ra*^fl/fl^ cells to CD127-negative *Ubc-cre*^ERT2^ *Il7ra*^fl/fl^ cells was calculated to account for CD127-positive escapees, favoritism for IL-7Rα-sufficient naïve phenotype CD8^+^ T cells became even more stark (**Figures 3D**). In contrast, for memory CD8^+^ T cells binding pooled gp276/D^b^/NP396/D^b^ tetramers or gp33/D^b^ tetramer, inducible loss of *Il7ra* again had only a relatively modest effect (**Figures 3E and S4A-S4C**). Furthermore, the proportion of CD127- positive, LCMV-specific escapee memory cells was relatively stable over time, in contrast to the strong selection observed for IL-7Rα-sufficient naïve CD8^+^ T cells (**Figures 3F, 3G, S4D, and S4E**). Tetramer-binding memory populations showed reduced proliferation in the absence of *Il7ra*, particularly for circulating populations (**Figures 3H, S4F, and S4G**). While trends were generally comparable to the P14 system, some differences in IL-7Rα dependence of proliferation were observed, particularly for the salivary gland (which exhibited greater IL-7Rα dependence in the chimera system). Among other explanations, it is possible that loss of IL-7Rα from half of the hematopoietic compartment in bone marrow chimeras resulted in a degree of lymphopenia, or alternatively that precursor frequency or TCR specificity may alter the degree of IL-7Rα dependence for proliferation. Taken together, resilience to loss of IL-7Rα was a conserved property across the P14 TCR transgenic and oligoclonal antigen-specific populations, with a conserved role for IL-7Rα in maintaining a normal proportion of proliferating memory CD8^+^ T cells. Loss of IL-7Rα did not result in rapid and severe attrition of CD8^+^ T cell memory.

### Efficient recall of memory CD8^+^ T cells can occur in the absence of IL-7Rα

While durability of CD8^+^ T cell memory was relatively normal after inducible deletion of *Il7ra*, capacity to recall in response to antigen might still be impaired. For example, IL-2Rα is required for normal programming of circulating memory CD8^+^ T cells during memory differentiation. In its absence, memory CD8^+^ T cell homeostasis is largely normal, with a severe defect appearing only upon recall^47^. Therefore, we deleted *Il7ra* from pre-formed memory P14 T cells, waited >4 weeks, and then subjected mice to heterologous challenge with *Listeria monocytogenes* expressing gp33 (Lm-gp33)^48^ (**Figure 4A**). Lm-gp33 was used to specifically recall gp33-specific P14 T cells and avoid confounding effects of antibodies against other LCMV antigens. To track the kinetics of the recall response, animals were bled over time. Irrespective of loss of IL-7Rα (and the subsequent 2-3 fold defect), in the blood we observed a robust increase in control and *Ubc-cre*^ERT2^ *Il7ra*^fl/fl^ P14 T cells after recall, near universal acquisition of Ki67 and granzyme B, and a profound shift from CD62L^+^ central memory phenotype cells to KLRG1^+^ effector phenotype cells (**Figures 4B-4H**). Therefore, loss of IL-7Rα during memory homeostasis did not obviously compromise the ability of circulating memory CD8^+^ T cells to recall.

**Figure 4:**
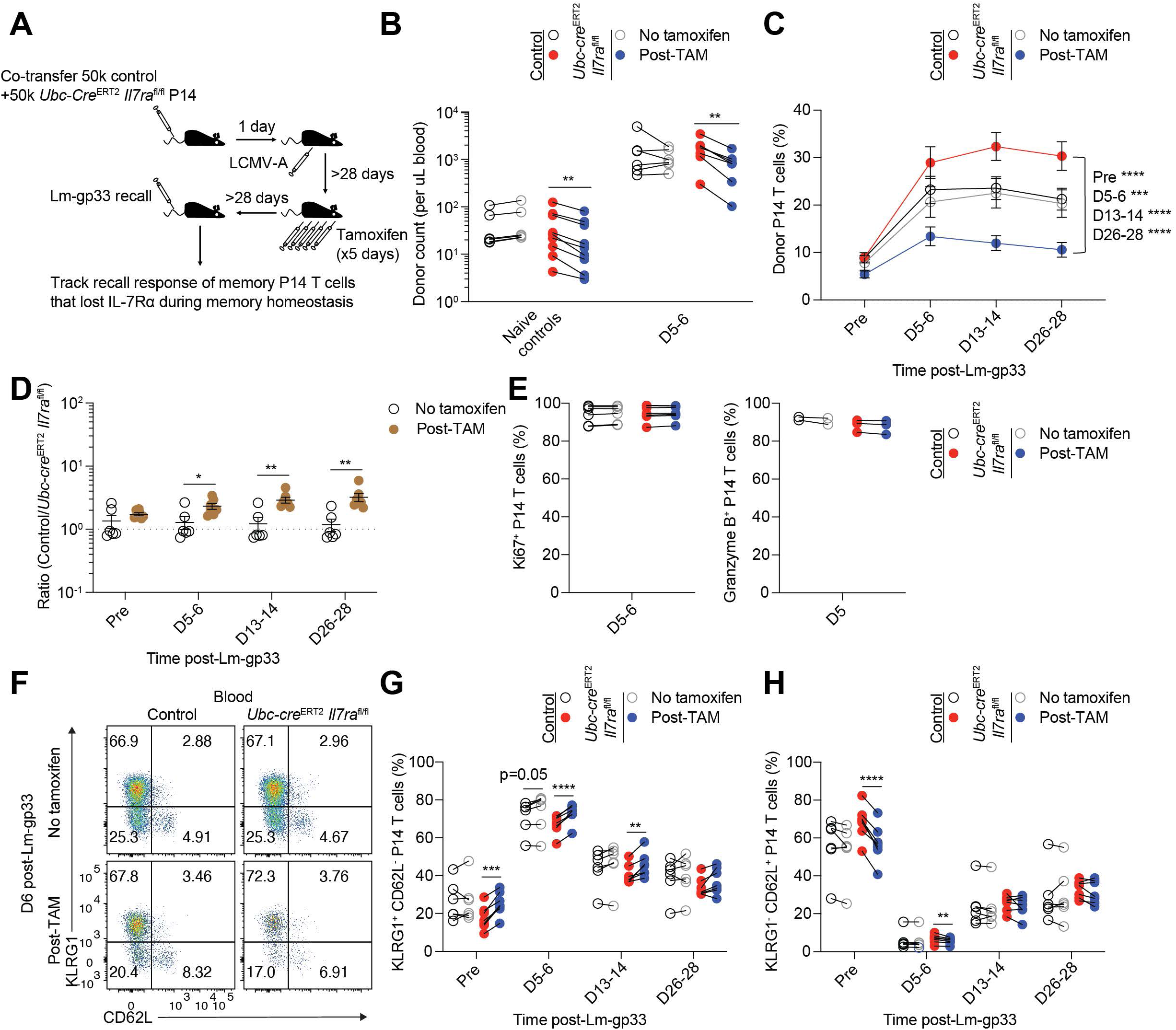
Ablation of the *Il7ra* during memory homeostasis does not significantly impair antigen-elicited recall of circulating memory. (A) Congenic control (*Ubc-cre*^ERT2^ *Il7ra*^+/+^ or *Il7ra*^fl/fl^) and *Ubc-cre*^ERT2^ *Il7ra*^fl/fl^ P14 T cells were co-transferred to recipients, followed by LCMV-Armstrong infection one day later. After resting to memory phase (>28 days), some mice were given i.p. tamoxifen for 5 consecutive days to initiate Cre-mediated recombination of the floxed *Il7ra* allele and rested (>28 days). Animals were then infected intravenously with ∼500,000 colony forming units of *Listeria monocytogenes* expressing gp33 (Lm-gp33) to reactivate P14 T cells. (B) Control and *Ubc-cre*^ERT2^ *Il7ra*^fl/fl^ P14 T cells were quantitated per µL blood from untreated and tamoxifen-treated Lm-gp33-reactivated mice and Lm-gp33-naïve controls. (C and D) Quantitation of the (C) percent (of CD8^+^ T cells) and (D) ratio of control and *Ubc-cre*^ERT2^ *Il7ra*^fl/fl^ P14 T cells from the blood of untreated and tamoxifen-treated Lm-gp33-reactivated mice. (E) Quantitation of the proportion of Ki67-expressing (left) and granzyme B-expressing (right) control and *Ubc-cre*^ERT2^ *Il7ra*^fl/fl^ P14 T cells from the blood of untreated and tamoxifen-treated Lm-gp33-reactivated mice. (F) Representative flow cytometry for KLRG1 and CD62L expression, (G) quantitation of the proportion of KLRG1^+^ CD62L^-^ P14 T cells as in (F), and (H) quantitation of the proportion of KLRG1^-^ CD62L^+^ P14 T cells as in (F) from control and *Ubc-cre*^ERT2^ *Il7ra*^fl/fl^ P14 T cells from untreated and tamoxifen-treated Lm-gp33- reactivated mice. Data are (B-E,G,H) compiled from 3 experiments (n=6-7/group), except granzyme B data (1 experiment, n=2-3/group), or (F) representative of 3 experiments (n=6-7/group). Error is expressed as ± S.E.M. Unpaired (D) and paired Student’s t tests (B-C,E,G,H). * p<0.05. ** p<0.01. *** p<0.001. **** p<0.0001.

### Deficiency in IL-15 signaling confers heightened dependence on IL-7Rα

While our data did not support the notion that IL-15-independent memory populations^28,29^ were more highly IL-7Rα-dependent, we considered the possibility that memory CD8^+^ T cells are instead regulated by the combination of IL-7 and IL-15 signaling. The lack of exclusive IL-15 dependence for some memory CD8^+^ T cell subsets does not preclude a role for IL-15 in concert with other cytokines in regulating such memory populations. As mentioned above, IL-15- independent memory CD8^+^ T cells respond robustly to IL-15 therapy^27^, supporting that memory cells adapt to γC cytokine availability irrespective of homeostatic requirements. Furthermore, enhanced severity of a combinatorial defect in IL-7 and IL-15 signaling has been proposed for circulating memory in the setting of a lymphopenic host or after transfer of day 10 effector cells^22,35,36,49^. Such redundancy could explain why IL-7 and IL-15 are individually less critical to CD8^+^ T cell memory than predicted and would have significant implications for their stability in the face of altered cytokine availability. We therefore considered whether IL-7Rα-deficient memory cells were more severely affected in the absence of IL-15.

To address this, control and *Ubc-cre*^ERT2^ *Il7ra*^fl/fl^ P14 T cells were transferred to IL-15- sufficient (*Il15*^+/+^ or *Il15*^+/-^) and *Il15*^-/-^ recipient mice followed by LCMV infection to elicit memory differentiation. First, by assessing control memory P14 T cell populations in IL-15-sufficient and - deficient recipients, we confirmed IL-15 independence of LCMV-elicited P14 memory cells in the IEL and FRT as previously reported^28^, while also establishing that liver memory cells have a moderate IL-15 dependence (**Figure S5A**). To determine the combinatorial effects of IL-7Rα/IL-15 deficiency, tamoxifen-treated IL-15-sufficient recipients were compared with untreated- and tamoxifen-treated *Il15*^-/-^ recipients. When tracking memory P14 populations in the blood after administration of tamoxifen, a greatly enhanced competitive advantage was observed for IL-7Rα- sufficient P14 cells in *Il15*^-/-^ recipients (**Figure 5A**). This led to profound selection for *Ubc-cre*^ERT2^ *Il7ra*^fl/fl^ CD127-expressing escapees of deletion over time; both features are consistent with a heightened requirement for IL-7Rα in the absence of IL-15 signaling (**Figure 5B**). When an adjusted ratio of control to genuinely CD127-negative *Ubc-cre*^ERT2^ *Il7ra*^fl/fl^ P14 cells was calculated, this revealed a nearly 100:1 competitive advantage for control cells in IL-15-deficient hosts (**Figure 5C**). In light of the outgrowth of CD127-expressing escapees of deletion, we took advantage of variation in the efficiency of *Il7ra* deletion using two independent cohorts, one with high efficiency deletion of *Il7ra* (Cohort 1, estimated at ∼90-95% for blood T_CM_ from concurrent wildtype recipients) and one with moderate efficiency deletion of *Il7ra* (Cohort 2, estimated at ∼70- 75% in the same manner) to address two related points (**Figures S5B and S5C**). In Cohort 1, escapees of deletion were quite rare, allowing clear assessment of the severe competitive disadvantage of lacking *Il7ra* across tissues and memory subsets in *Il15*^-/-^ recipients by ratio (**Figures 5D-5F and S5D**). However, it was challenging to accurately assess the phenotype of the remaining *Ubc-cre*^ERT2^ *Il7ra*^fl/fl^ P14 cells, especially in NLT, because of their extreme scarcity, though it appeared that a significant majority of these cells were CD127-positive escapees of Cre-mediated recombination (**Figure 5E, 5F, and S5E**). However, in Cohort 2, while the higher proportion of escapees limited the ratio that could be observed between donors (**Figure S6A**), it allowed observation of competition within the *Ubc-cre*^ERT2^ *Il7ra*^fl/fl^ P14 population after tamoxifen. While CD127-positive escapees in B6 recipients had only a small advantage, persisting *Ubc- cre*^ERT2^ *Il7ra*^fl/fl^ P14 cells in *Il15*^-/-^ hosts became nearly 100% CD127-positive in circulating populations and exhibited nearly normal gMFI for CD127 in lymphoid and NLT sites, consistent with selection for IL-7Rα-sufficient escapees (**Figures 5G and S6B-S6E**). Across both cohorts, combined deficiency in IL-7Rα and IL-15 resulted in reductions in P14 counts (**Figure S6F**). Therefore, lack of IL-15 imposed a heightened requirement for IL-7Rα and revealed cooperativity between γC cytokines which conferred resilience to altered cytokine availability.

**Figure 5:**
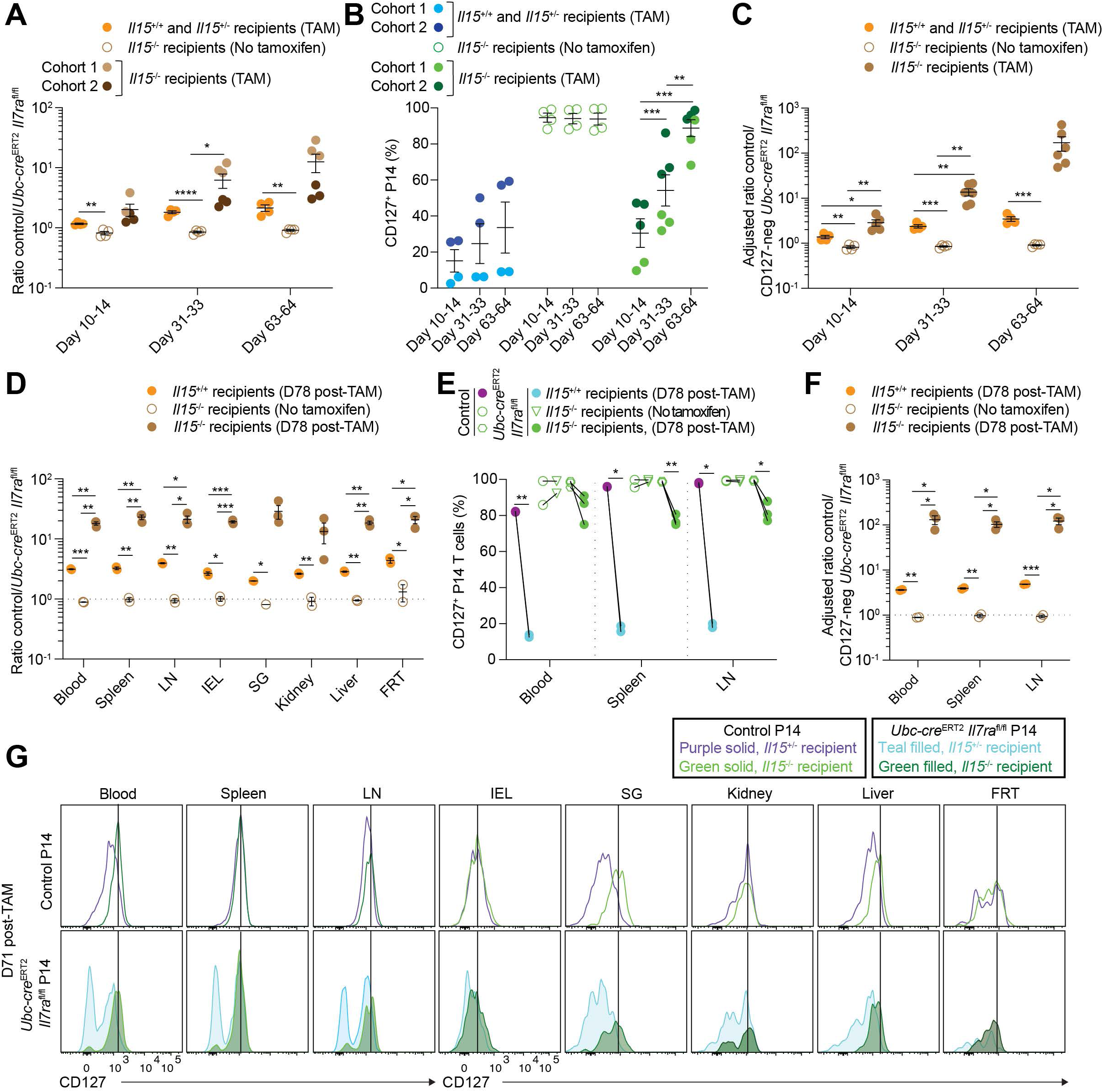
Redundancy between IL-7 and IL-15 affords resilience to loss of individual cytokines. Congenic control (*Ubc-cre*^ERT2^ *Il7ra*^+/+^ or *Il7ra*^fl/fl^) and *Ubc-cre*^ERT2^ *Il7ra*^fl/fl^ P14 T cells were co-transferred to IL-15-sufficient (*Il15*^+/+^ or *Il15*^+/-^) or -deficient recipients, followed by LCMV- Armstrong infection one day later. After resting to memory phase (>28 days), some mice were given i.p. tamoxifen for 5 consecutive days to initiate Cre-mediated recombination of the floxed *Il7ra* allele and tracked over time in the blood via gp33/D^b^ tetramer staining and congenic markers. (A-C) Cohort 1 and 2 control and *Ubc-cre*^ERT2^ *Il7ra*^fl/fl^ memory P14 T cells were quantitated for the (A) ratio, (B) proportion CD127^+^, and (C) adjusted ratio of control to CD127-negative *Ubc-cre*^ERT2^ *Il7ra*^fl/fl^ memory P14 T cells from the blood of untreated and tamoxifen-treated mice. (D-F) Cohort 1 control and *Ubc-cre*^ERT2^ *Il7ra*^fl/fl^ P14 T cells were quantitated for the (D) ratio, (E) proportion CD127^+^, and (F) adjusted ratio of control and CD127-negative *Ubc-cre*^ERT2^ *Il7ra*^fl/fl^ memory P14 T cells from untreated and tamoxifen-treated mice. (G) Representative flow cytometry for CD127 expression on Cohort 2 control and *Ubc-cre*^ERT2^ *Il7ra*^fl/fl^ P14 T cells from *Il15*^+/-^ or *Il15*^-/-^ recipients. Data are (A-C) compiled from 2 experiments (3-6/group) or (D-G) representative of/derived from 1 experiment (n=2-3/group, except SG, No tamoxifen [n=1]). Error is expressed as ± S.E.M. Unpaired (A-D,F) or paired (E) Student’s t tests. * p<0.05. ** p<0.01. *** p<0.001. **** p<0.0001. See also Figures S5 and S6.

### Integrated regulation of IL-7 and IL-15 signaling modulates CD8^+^ T cell memory

Previous work on predominantly naïve CD8^+^ T cells revealed that IL-7 (and other cytokines including IL-15) downregulated mRNA and protein for IL-7Rα as a mechanism to conserve IL-7 and allow signaling for as many cells as possible^42^. However, it has not been determined whether a similar mechanism holds for memory cells. CD127 expression appeared to be upregulated on control P14 T cells in *Il15*^-/-^ recipient mice (**Figure 5G**), so we quantitated this across IL-15- sufficient and -deficient animals. Strikingly, CD127 was upregulated on control memory P14 T cells from most tissues of *Il15*^-/-^ recipients (**Figures 6A and 6B**). When CD127 levels were assessed on circulating memory CD8^+^ T cells after in vivo IL-7 or IL-15 complex treatment (IL-7c and IL-15c, a more potent and long-lasting method to administer cytokine^27,50–52^), in both cases CD127 was downregulated (**Figures 6C-6F**). Downregulation of IL-7Rα by IL-15c treatment suggested that the observed increase in CD127 staining across memory subsets in *Il15*^-/-^ recipients could be due to a lack of IL-15 signaling and may act as a compensatory mechanism to allow enhanced IL-7 sensitivity. As IL-7Rα was regulated by IL-15 signaling, we then asked whether loss of IL-7Rα from homeostatic memory P14 T cells altered IL-15 responsiveness. We first assessed whether inducible deletion of *Il7ra* resulted in upregulation of IL-2Rβ (CD122). However, there was no clear evidence of this (**Figures S7A and S7B**), suggesting that any altered sensitivity of IL-7Rα-deficient memory CD8^+^ T cells to IL-15 did not occur at the level of receptor expression. We next treated mice with a low dose of IL-15 (2 µg) to induce an intermediate response^53^ of memory CD8^+^ T cells and thereby assess whether IL-7Rα-sufficient and -deficient memory cells differed in IL-15 sensitivity. Indeed, loss of IL-7Rα from homeostatic memory cells impaired proliferation in response to low dose IL-15 for circulating memory P14 T cells as well as some resident populations to a more modest extent (**Figures 6G and S7C**). This phenotype was enhanced when specifically gating on CD127-negative *Ubc-cre*^ERT2^ *Il7ra*^fl/fl^ P14 T cells in the blood and lymphoid tissues (**Figure 6H**). Taken together, IL-7 and IL-15 signaling exhibit cross-regulation, suggesting both compensatory and cooperative relationships between these cytokines to support memory CD8^+^ T cells

**Figure 6:**
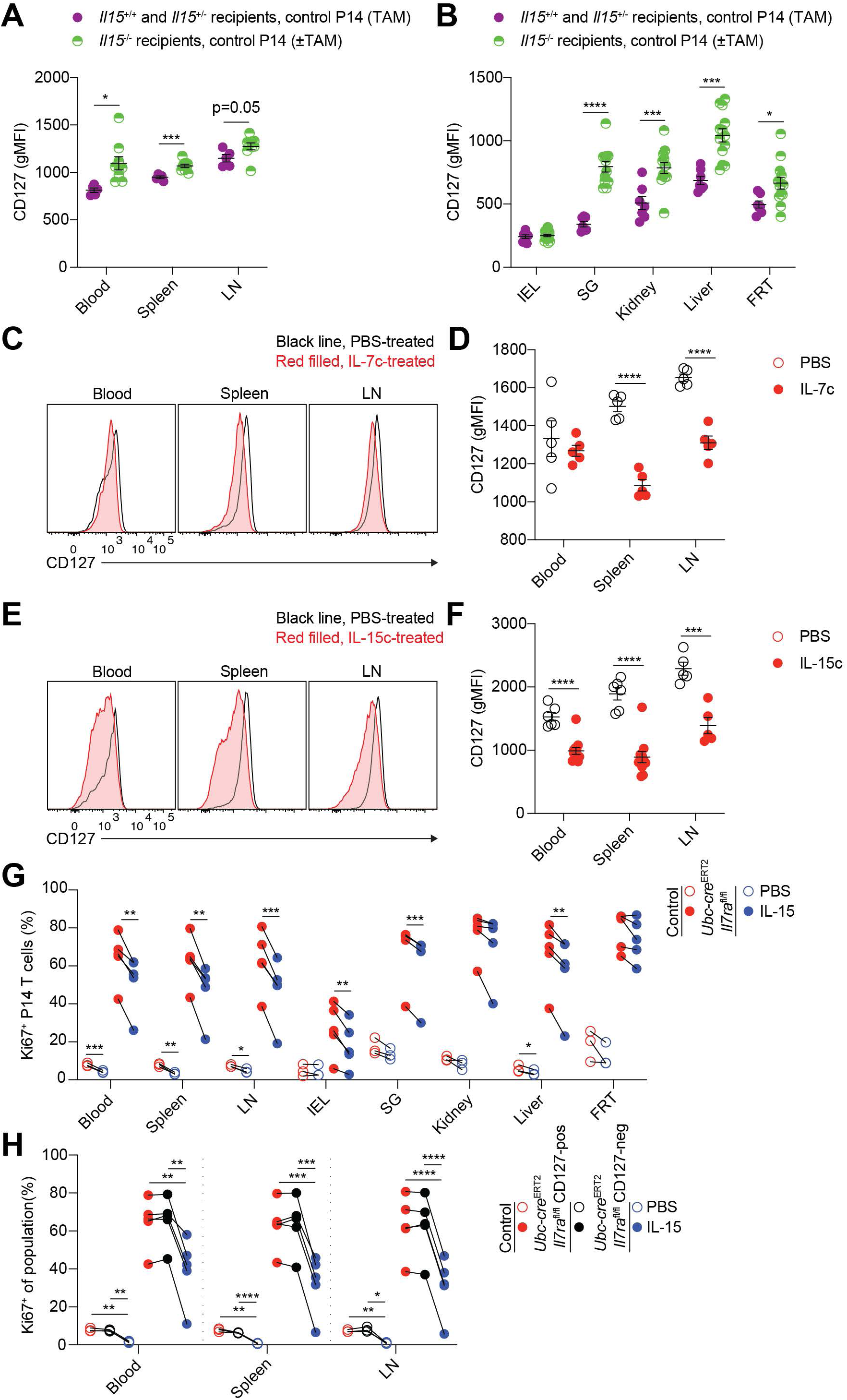
Cross-regulation of IL-7 and IL-15 signaling modulates CD8^+^ T cell memory. (A and B) As in Figure 5, congenic control (*Ubc-cre*^ERT2^ *Il7ra*^+/+^ or *Il7ra*^fl/fl^) and *Ubc-cre*^ERT2^ *Il7ra*^fl/fl^ P14 T cells were co-transferred to IL-15-sufficient (*Il15*^+/+^ or *Il15*^+/-^) or -deficient recipients, followed by LCMV-Armstrong infection one day later. After resting to memory phase (>28 days), some mice were given i.p. tamoxifen for 5 consecutive days to initiate Cre-mediated recombination of the floxed *Il7ra* allele and tracked over time in the blood via gp33/D^b^ tetramer staining and congenic markers. Quantitation of CD127 gMFI for (A) lymphoid tissue- and (B) non-lymphoid tissue-localized control memory P14 T cells from untreated and tamoxifen-treated mice (some data points are also shown in Figs. S5E, S6D, and S6E). (C-F) Congenic wildtype P14 T cells were transferred to recipients, followed by LCMV-Armstrong infection one day later. After resting to memory phase (>28 days), mice were given PBS and (C and D) IL-7c or (E and F) IL-15c on day 0 and 2 with sacrifice on day 4. (C and E) Representative flow cytometry for CD127 expression on memory P14 T cells from PBS- and cytokine complex-treated mice and (D and F) quantitation of CD127 gMFI as in (C and E). (G, H) Congenic control (*Il7ra*^fl/fl^) and *Ubc-cre*^ERT2^ *Il7ra*^fl/fl^ P14 T cells were co-transferred to recipients, followed by LCMV-Armstrong infection one day later. After resting to memory phase (>28 days), all mice were given i.p. tamoxifen for 5 consecutive days to initiate Cre-mediated recombination of the floxed *Il7ra* allele and then rested (>28 days). Mice were then treated with PBS or 2 µg IL-15. (G) Quantitation of the proportion of Ki67-expressing control and *Ubc-cre*^ERT2^ *Il7ra*^fl/fl^ memory P14 T cells from PBS or IL-15-treated mice. (H) Quantitation of the proportion of Ki67-expressing control, *Ubc-cre*^ERT2^ *Il7ra*^fl/fl^ CD127^+^, and *Ubc- cre*^ERT2^ *Il7ra*^fl/fl^ CD127^-^ memory P14 T cells from PBS or IL-15-treated mice. Data are (A,B,D,F-H) compiled from 2-3 experiments (n=3-13/group) or (C,E) representative of 2-3 experiments (n=4- 12/group). Error is expressed as ± S.E.M. Unpaired (A,B,D,F) or paired (G,H) Student’s t tests. * p<0.05. ** p<0.01. *** p<0.001. **** p<0.0001. See also Figure S6.

## DISCUSSION

The dogma that IL-7 and IL-15 each fulfill exclusive roles essential for memory CD8^+^ T cell maintenance has been widely accepted in the field, despite repeated findings inconsistent with this model^22,28,29,38,41^. Previous experimental approaches to address the role of IL-7 signaling are difficult to interpret because of potential confounding effects on hematopoiesis, T cell development, naïve T cell homeostasis, and memory differentiation, with substantial T-lymphopenia when IL-7 or its receptor are ablated or blocked. Using inducible deletion of a floxed *Il7ra* allele, we rigorously assessed the cell-intrinsic role of IL-7Rα in homeostasis of defined, antigen-specific populations of memory CD8^+^ T cells in lymphoreplete hosts. Counter to prevailing models, IL-7Rα played a nuanced role in maintenance of memory CD8^+^ T cells, with ablation leading to impaired proliferation of memory CD8^+^ T cells in most sites and gradual attrition. However, loss of IL-15 greatly augmented dependence on IL-7Rα, even among T_RM_ populations that are classified as IL-15-independent. These data, together with the observation that loss of either IL-7 or IL-15 signaling affected the signaling pathway for the other cytokine, argue that memory CD8^+^ T cells dynamically adapt to available homeostatic cytokines, and thus that IL-7 or IL-15 “dependence” is not fixed.

A major focus of this study was to investigate the roles of IL-7 and IL-15 in T_RM_. Because these populations adapt to the tissues in which they reside^6–12^, it would not be surprising if T_RM_ in distinct sites exhibited different homeostatic requirements, consistent with reduced IL-2Rβ and IL-7Rα expression on some T_RM_^26,27^. Indeed, previous reports^28,29^ indicated T_RM_ populations in some sites were IL-15-independent, while T_RM_ in other tissues stringently required IL-15 much like circulating memory, as confirmed in our study. It was unclear whether IL-15-independent T_RM_ would exhibit heightened sensitivity to IL-7 or, alternatively, no requirement for either IL-7 or IL-15 at all. Instead, induced *Il7ra* ablation caused gradual decline of memory CD8^+^ T cells in all sites, a resilience that is presumably acquired during memory differentiation as it is not shared with naïve CD8^+^ T cells, as previously reported^44^. Interestingly, the most notable impact of IL-7Rα loss was impaired basal proliferation of most memory CD8^+^ T cells (although this was muted among T_RM_ in some sites), in contrast to the prevailing notion that IL-7 predominantly regulates memory cell survival in normal (i.e. non-irradiated) hosts^20,22^. Goldrath et al previously noted that blockade of IL-7Rα might elicit lymphopenia-induced proliferation (i.e. via IL-15). This likely confounded previous analyses, but using inducible deletion of *Il7ra*, we now describe a role for IL-7Rα in basal proliferation of most memory CD8^+^ T cell subsets. While changes were not observed in cell death, loss of IL-7Rα during memory homeostasis did reduce Bcl2 levels on circulating memory cells, as predicted based on previous work^20,39,40^. Therefore, future studies should take into account the impact of IL-7Rα on both self-renewal and survival of memory CD8^+^ T cells. Despite their defect in basal proliferation, IL-7Rα-deficient circulating memory CD8^+^ T cells were capable of robust antigen-elicited proliferation during recall. This suggests that cells which are quiescent prior to TCR engagement are not excluded from recall in favor of memory cells actively undergoing basal proliferation.

While loss of IL-7Rα did perturb basal proliferation, memory CD8^+^ T cell subsets experienced only gradual attrition over time (2-3 fold up to 2.5 months after tamoxifen), likely due to low requirements for self-renewal. Our findings appear to stand in contrast to previous studies of CD8^+^ T cells lacking normal IL-7Rα signaling during activation and memory differentiation, which showed a competitive defect in excess of 10:1 approximately one month after transfer^20,39^. However, our study focused on the role of IL-7Ra in mediating homeostasis of pre-formed memory CD8^+^ T cells, not the differentiation of such cells. Indeed, a role for IL-7Rα has previously been shown in supporting the generation of CD8^+^ memory-precursors and establishing a durable memory CD8^+^ T cell pool^35,36,39,41^. By inducibly ablating *Il7ra* after memory CD8^+^ T cell differentiation, we could selectively interrogate the contribution of IL-7Rα to memory CD8^+^ T cell homeostasis. The capacity of memory CD8^+^ T cells to persist after loss of IL-7 signaling may be critical for their stability during temporary alterations in IL-7 availability in vivo. This could potentially occur during inflammation or tissue damage affecting IL-7-producing stromal cells, though little is known regarding physiological perturbations in IL-7 expression^20,54–57^.

Our data also indicated that CD8^+^ T cell memory populations in an IL-15-deficient environment adopted a heightened reliance on IL-7 signaling, likely resolving the enigma of IL-15-independent memory subsets. Furthermore, we demonstrate in vivo crosstalk between IL-7 and IL-15 signaling in memory CD8^+^ T cells. Previous studies of naïve T cell populations showed that IL-7Rα expression is reduced by stimulation with diverse cytokines (including IL-7 itself and IL-15)^42^. These findings were confirmed in vivo for memory CD8^+^ T cells, and we also showed that surface levels of IL-7Rα are enhanced on memory P14 T cells in *Il15*^-/-^ hosts. As downregulation of IL-7Rα has previously been proposed as an IL-7 conservation mechanism to allow efficient distribution of IL-7 amongst a population of cells^42^, it may be that the previously reported defects in CD8^+^ T cell memory in *Il15*^-/-^ mice^19,21–25,28,29^ are not solely due to IL-15 deficiency, but also to inefficient use of IL-7 at the population level. In turn, the modest defect we describe after loss of IL-7Rα from homeostatic memory cells may not be exclusively due to loss of IL-7 sensitivity. We found that loss of IL-7Rα can impair proliferation of memory CD8^+^ T cells in response to IL-15, suggesting that intact IL-7 signaling also has a secondary effect on IL-15 responsiveness. In a physiological context, these mechanisms could potentially conserve IL-7 while maximizing IL-7 and IL-15 signal strength across a memory population, thereby moderating the effect of local changes in cytokine availability due to consumption and competition. We propose that IL-7 and IL-15 collaborate to support memory CD8^+^ T cells in homeostasis, not only via redundancy (as both strongly activate STAT5 as well as other shared pathways, including PI3K/Akt and MAP kinases), but also by cross-regulation. This affords a measure of resilience and adaptability in the absence of signaling for either cytokine, but results in a severe defect across homeostatic memory populations when both signals are absent. During inflammation, cytokine availability undergoes major alterations, including marked increases in IL-15 during viral infection due to type 1 interferon^53^ and also altered cytokine turnover. The flexibility of memory CD8^+^ T cells to adapt to individual loss (or gain) of IL-7 and IL-15 signaling may help to ensure their stability during inflammation. It is intriguing to speculate that in an inflammatory setting, memory CD8^+^ T cells may rely on an expanded cadre of cytokines, with important implications for their stability and long-term maintenance.

## LIMITATIONS OF THE STUDY

The present study (and many previous publications) employed memory CD8^+^ T cells which differentiated in *Il15*^-/-^ recipient mice and therefore lacked IL-15 throughout both differentiation and memory homeostasis, which may overrepresent the requirement for IL-15 during memory homeostasis. Future work should address the specific role for IL-15 signaling in memory CD8^+^ T cell homeostasis. The mechanism by which IL-7Rα licenses full IL-15 responsiveness is not clear. Preliminary data suggest that strong IL-15 signals (i.e. IL-15 complexes) can overcome the requirement for homeostatic IL-7Rα, suggesting that IL-7Rα acts to enhance the response to IL-15 at more physiological levels.

## ACKNOWLEDGEMENTS

N.N.J. is a Damon Runyon Fellow supported by the Damon Runyon Cancer Research Foundation (Grant No. DRG-2427-21) and the National Institute of Allergy and Infectious Diseases (Grant No. 1K22AI177360-01). K.M.W. was supported by National Cancer Institute (NCI) (Grant No. F30 CA250321). N.J.M. was supported by NCI (Grant No. K00 CA245735). K.M.A was supported by NIAID (Grant No. T32 AI007313). This work was funded by NIAID Grant No. R01 AI038903 to S.C.J. The authors thank M. Jenkins and K. Osum for critical reading of the manuscript. The authors acknowledge the S.C.J., D.M., K. A. Hogquist, and M. Jenkins laboratories for valuable feedback and reagents. The gp33/D^b^, gp276/D^b^, and NP396/D^b^ monomers were obtained through the NIH Tetramer Core Facility. The authors dedicate this work to the memory of Leo Lefrancois and Charlie Surh.

## AUTHOR CONTRIBUTIONS

Conceptualization: N.N.J., S.C.J. Methodology: N.N.J., T.S.D., C.P., T.A.D., K.M.A., D.M, S.C.J. Formal analysis: N.N.J., S.C.J. Investigation: N.N.J., T.S.D., N.J.M, K.M.W., C.P., T.A.D., K.E.B., W.J.V., K.M.A. Resources: D.M., S.C.J. Data Curation: N.N.J., S.C.J. Writing-Original Draft: N.N.J., S.C.J. Writing-Review & Editing: All Authors. Visualization: N.N.J., S.C.J. Supervision: N.N.J., S.C.J.

## KEY RESOURCES TABLE

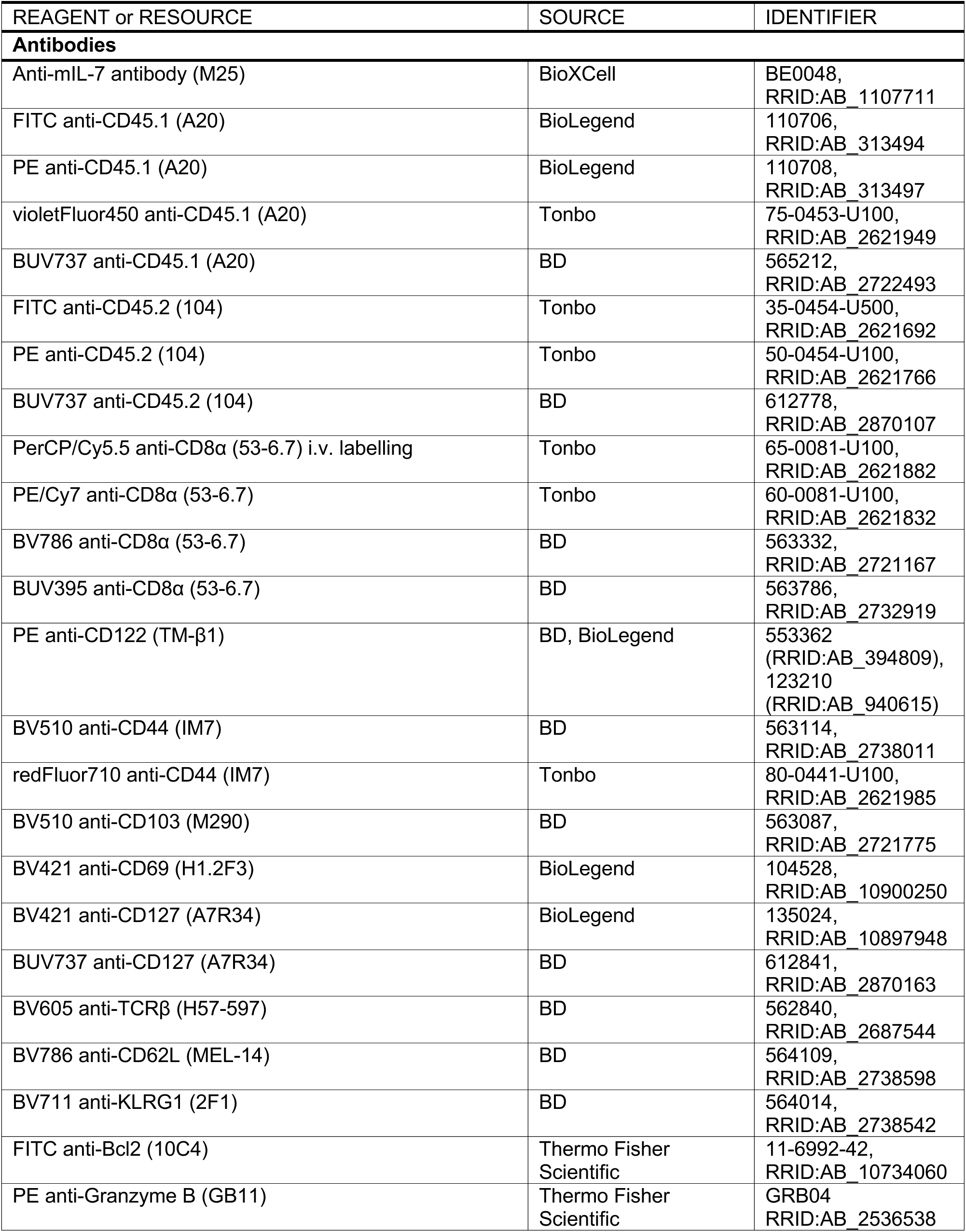
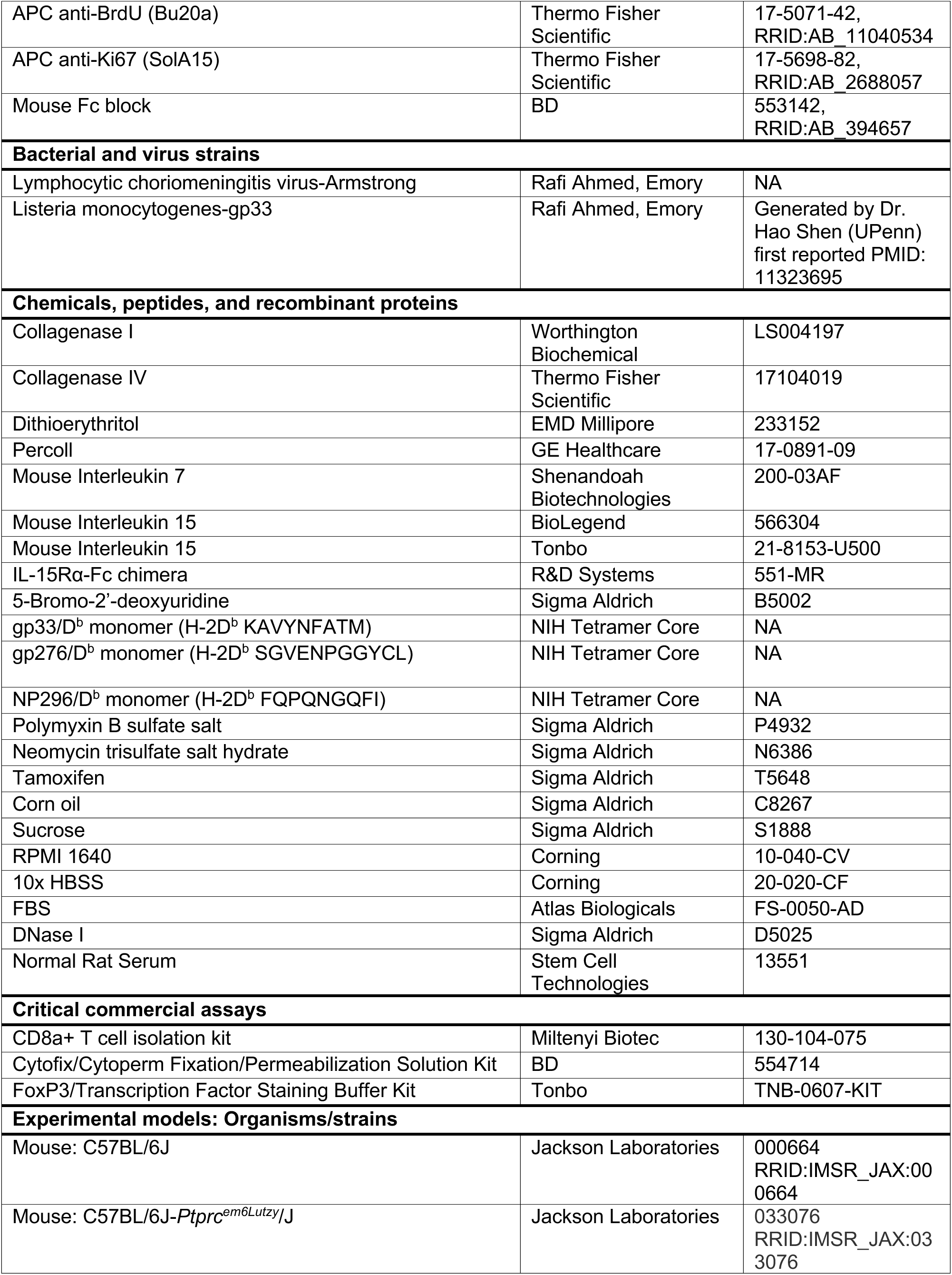
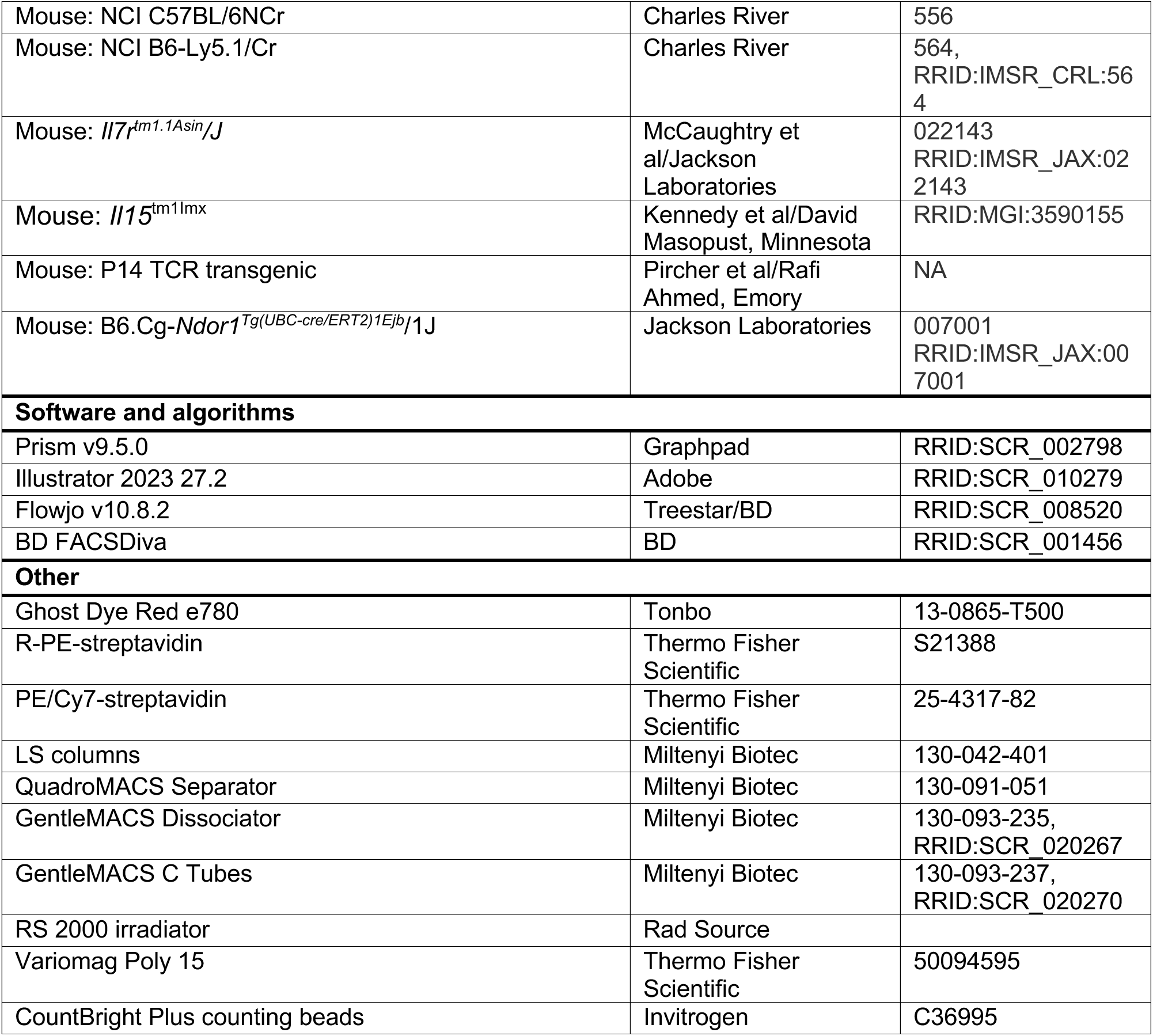

## RESOURCE AVAILABILITY

### Lead contact

Further information and requests for resources and reagents should be directed to the lead contact, Stephen Jameson.

### Materials availability

This study did not generate unique reagents.

## EXPERIMENTAL MODEL AND STUDY PARTICIPANT DETAILS

### Mice

See also the Key Resources Table. C57BL/6J and JAXboy mice were purchased from Jackson Laboratory for use as *Il7ra*^fl/fl^ P14 transfer recipients. *Il15*^-/-^ mice^19^ were backcrossed to C57BL/6J and bred in house as heterozygous by heterozygous and heterozygous by homozygous for use as *Il7ra*^fl/fl^ P14 transfer recipients; in some experiments C57BL/6J were also used as *Il15*^+/+^ recipients. For all lines backcrossed, speed congenics from Transnetyx were used. CD45.1/.1 and CD45.2/.2 C57BL/6N (purchased from Charles River) were used as recipients for mixed bone marrow chimera experiments and cytokine complex therapy experiments. *Il7ra*^fl/fl^ mice^58^ with a floxed exon 3 of the *Il7ra* gene were purchased from Jackson Laboratory on a mixed background (used for BM chimera experiments) and were backcrossed 5-6 generations to C57BL/6J (used for adoptive transfer experiments). *Ubc-cre*^ERT2^ mice were purchased from Jackson Laboratory. LCMV-gp33/D^b^-specific P14 TCR transgenic mice^59^ were backcrossed and maintained on the C57BL/6J and C57BL/6N backgrounds as separate lines. C57BL/6J P14 mice were intercrossed with *Il7ra*^fl/fl^, *Ubc-cre*^ERT2^, and Jaxboy mice to generate congenically distinct donor mice. *Ubc- cre*^ERT2^-positive male breeders were crossed to female *Il7ra*^fl/fl^ mice lacking the Cre transgene and offspring were typed for the floxed allele and the recombined version to identify spontaneous germline deletion in the absence of tamoxifen^60^, which was rare for this strain. Donors (control and experimental derived from the same breeding colony, sibling/littermates used when possible) and recipients were used between 6-14 weeks of age. Female animals were used as recipients (and donors) in our study, due to limitations on housing males for long-term memory experiments due to aggression. Animals were housed under specific pathogen-free conditions at the University of Minnesota; infections were performed in a BSL2 animal facility. All animal procedures were approved by the institutional animal care and use committee of the University of Minnesota.

### Pathogens and infections

Infections and treatments: Mice were infected intraperitoneally (i.p.) with LCMV Armstrong at 2*10^5^ PFU. Mice were rechallenged intravenously with ∼500,000 colony forming units of Listeria monocytogenes expressing gp33^48^.

## METHOD DETAILS

### Tamoxifen treatment

Tamoxifen was dissolved in corn oil at 20 mg/mL for >5 hours at 37C with 250 rpm shaking (New Brunswick Scientific, Excella E24 Incubator Shaker Series) and vortexing, with protection from light. Tamoxifen in oil was prepared fresh for each course of treatment. Mice were given 100 uL (2 mg) i.p. on 5 consecutive days, per the Jackson Laboratory (https://www.jax.org/research-and-faculty/resources/cre-repository/tamoxifen). For P14 cohorts 1 and 2, high efficiency or moderate efficiency deletion was confirmed respectively to be conserved across different batches of tamoxifen and for both naïve and memory cells derived from the same donor. Variability in Cre activity, even between littermates, has been previously reported^60^.

### BrdU treatment

1 mg was given i.p. at the start of treatment, with continuous BrdU administration via drinking water ad libitum (0.8 mg/mL in 2% sucrose water with protection from light). BrdU water was changed every other day.

### Cytokine treatment

For low dose IL-15 treatment, 2 μg mIL-15 were given i.p. as published^53^ on days 0 and 2 before harvest on day 3. For IL-7 and IL-15 complex treatment, mice were treated on days 0 and 2 before harvest on day 4. IL-7c were formed with 1.5 μg mIL-7 and 7.5 μg anti-IL-7 antibody per mouse, combined in equal volume and incubated for 2-3 minutes at RT, followed by dilution in PBS to 200 uL, and were placed on ice before injection i.p.^52^. IL-15c were prepared as previously described^27,61^.

### CD8^+^ T cell adoptive transfers

CD8^+^ T cells were isolated from the spleens of donor P14 TCR transgenic mice with the Miltenyi Biotec CD8a^+^ T cell isolation kit, mouse and Miltenyi Biotec LS columns. 50,000 cells from each donor were transferred to naive recipient mice by retro-orbital or tail vein injection before LCMV-A infection to elicit memory. To track naïve P14 T cells, ∼1.5-2M P14 T cells from each donor were co-transferred, followed by tamoxifen treatment as above within a week (generally ∼4 days post-transfer).

### Mixed BM chimeras

Recipient mice were given split-dose irradiation (2 doses of 500 rads, RS 2000 irradiator [Rad Source]) on two consecutive days, followed by intravenous transfer of 4M BM cells from each donor (isolated from hind limb femurs and tibias without red blood cell lysis). Mice were given two weeks of antibiotic-treated water (polymyxin B sulfate salt 15 mg/L, neomycin trisulfate salt hydrate 40 mg/L) during reconstitution. After >8 weeks, animals were infected with LCMV- Armstrong as above and rested for >4 weeks to establish memory. Animals were then treated with tamoxifen as above.

### Tissue harvests

5 minutes before harvest, mice were given 3 μg anti-CD8α PerCP/Cy5.5 by retro-orbital injection^45^. Animals were then cheek bled into heparin and sacrificed for tissue harvest. The inguinal LNs and the spleen were collected into harvest media (either RPMI 1640 supplemented with 5% fetal bovine serum [heat inactivated before use] or 1x Hanks’ balanced salt solution [HBSS] supplemented with 2.38 g/L Hepes], 2.1 g/L sodium bicarbonate, and 5% FBS) and passed through a 70-μm cell strainer. For IEL, the small intestine (SI) was excised and divested of fat and doused in IEL media (1x HBSS supplemented with Hepes, sodium bicarbonate, and 2% FBS) to keep moist. The Peyer’s patches were then removed and fecal contents were extruded from the lumen, which was then cut open. The sample was cut into 3-4 longitudinal sections, vortexed, and left on ice in 20 mL IEL medium. IEL medium was decanted, and tissue was washed with 30 mL of fresh IEL medium. Tissue was then transferred to 50 mL Erlenmeyer flasks with stir bars and 30 mL of IEL dithioerythritol (DTE) media supplemented with additional FBS to 5% and 154 mg/L dithioerythritol. After 30 min of stirring at 37 °C on a Variomag Poly 15, the supernatant was decanted through a 70 μm filter. 20 mL of IEL DTE media was added, followed by vortexing. After allowing tissue to settle, the supernatant was filtered and combined with the previous supernatant fraction. For liver, the organ was excised, avoiding the gallbladder, and placed in 5 mL harvest media on ice. It was then disrupted using a GentleMACS C tube on a GentleMACS Dissociator (m_spleen_01.01 twice) and filtered. For SG, kidney, FRT. SGs (submandibular, part of the sublingual) were excised, and cervical LNs were removed if present. The parotid was excluded to avoid lymphoid contamination. The kidney capsule was removed during isolation. The FRT was collected inclusive of the ovaries to the vagina and was bisected open. All tissues were placed in harvest media on ice. Tissues were finely minced with scissors and transferred to Erlenmeyer flasks with stir bars. 30 mL of collagenase solution (RPMI 1640 supplemented with 1 mM MgCl2, 1 mM CaCl2, 111.6 mg/L Hepes, 292 mg/L L-glutamine, and 5% FBS) containing 0.364 mg/mL collagenase I (SG, kidney) or 0.5 mg/mL collagenase IV (FRT) was added. Tissues were then incubated at 37 °C with stirring for 45-55 min (SG, kidney) or 60-70 min (FRT). After digestion, supernatants were filtered, and the remaining tissue was transferred to a GentleMACS C tube as above for liver. GentleMACS contents were then filtered and combined with the previous fraction. All NLTs were pelleted and resuspended in 5 mL of room temperature (RT) 44% Percoll (diluted with RPMI 1640), before underlay with 3 mL of RT 67% Percoll (diluted with PBS). Percoll was mixed with 10x PBS before use. Samples were centrifuged for 20 min at 800 g at RT, with minimum acceleration and deceleration. The interface was collected, diluted with harvest media, and used for downstream analysis. Red blood cells in blood and spleen samples were lysed with ACK buffer (150 mM ammonium chloride, 1 mM potassium bicarbonate, and 0.1 mM EDTA [ethylenediaminetetraacetic acid] in water) before staining.

### Flow Cytometry

Samples and splenic single stain controls were washed in fluorescence-activated cell sorter (FACS) buffer (2% FBS, 2 mM EDTA in 1x PBS), followed by Fc blocking for 5 min. Antibodies/viability dye for staining were then added for 20 min at 4 °C with concurrent tetramer staining when used. Samples were then washed before fixation, washed after fixation, and stored in FACS buffer at 4 °C before intracellular staining 1 to 2 d later. Antibodies/viability dyes are listed in the Key Resources Table. All surface antibodies were used at 1/200, except anti-CD69 and anti-CD127 (1/100). Viability dye was used at 1/1,000. Intracellular Staining. For Ki67 (1/200), Bcl2 (1/50), and Granzyme B staining (1/200), the Tonbo Foxp3/Transcription Factor Staining Buffer Kit was used according to the standard protocol, with an additional 15-min incubation in 1x Perm buffer plus 2% Normal Rat Serum before antibody staining for 45-60 min at RT. BrdU staining was done using a modified protocol for the BD Cytofix/Cytoperm Fixation/Permeabilization Solution Kit. Briefly, cells were washed in 1x Perm/Wash (P/W) buffer and then incubated for 10 min at 4 °C in 1x P/W buffer plus dimethyl sulfoxide (1 part 10x P/W, 1 part DMSO, and 8 parts water). Samples were then washed in 1x P/W buffer and refixed in Cytofix/Cytoperm buffer for 5 min at RT, followed by washing in 1x P/W buffer. Cells were then digested with deoxyribonuclease (DNase) I at 37 °C for at least 50 min. DNase was stored at -80 °C as a 2 mg/mL stock in PBS and diluted to 600 μg/mL in 1x PBS immediately before use. After digestion, cells were washed in 1x P/W buffer and stained with anti-BrdU antibody (1/100) for 50-60 min at RT. Cells were washed with 1x P/W buffer and then with FACS buffer. All NLT samples were filtered, and CountBright Plus counting beads were added to flow tubes before analysis. Samples were acquired using BD LSRII, LSRFortessa X20 and X30, and LSRFortessa flow cytometers and FACSDiva software. Analysis was done using FlowJo v7 and v10. Singlet lymphocytes were gated by forward scatter area (FSC-A)/side scatter area (SSC-A) and FSC-A/FSC-width (FSC-W). Live cells were then gated according to i.v. labeling status (blood, i.v.-positive; LN, IEL, SG, FRT, i.v.-negative; spleen and kidney, i.v.-low; liver, not gated by i.v. labeling status). CD8α^+^/TCRβ^+^ cells were then divided by CD45 congenic staining into host and donor populations, with gp33/D^b^ tetramer staining to exclude the host for *Il15*^-/-^ and naïve CD8^+^ T cell experiments. Tetramer binding cells were gated as tet^+^ CD44^hi^. To validate BrdU gating, each experiment included a control animal not given BrdU, tissues from which were processed and stained alongside the other samples to serve as negative controls. For subsetting of memory, blood, spleen, and LN P14 T cells were gated as KLRG1^+^ CD62L^-^ (long-lived effector cells), KRG1^-^ CD62L^-^ (T effector memory), and KLRG1^-^ CD62L^+^ (T central memory). For the adjusted ratio of control to CD127-negative *Ubc-cre*^ERT2^ *Il7ra*^fl/fl^ P14 T cells, adjusted ratios were only calculated for recipients that had received tamoxifen and could therefore have lost the IL-7Rα. For animals not receiving tamoxifen, the simple ratio of donor to donor was used, meaning the ratios shown for no tamoxifen recipients are identical for the simple and adjusted ratio graphs. For salivary gland, P14 T cells were split into two CD69^+^ resident subsets, differing in expression of CD103. In rare cases, dominance of CD69/CD103 double-negative cells in SG samples revealed lymphoid tissue contamination, and these samples were excluded. For kidney and liver, P14 T cells were split into CD69^+^ resident and CD69^-^ circulating memory.

### Tetramer reagents

Tetramers were made using biotinylated monomers (H-2D^b^ KAVYNFATM [gp33/D^b^]; H-2D^b^ SGVENPGGYCL [gp276/D^b^]; H-2D^b^ FQPQNGQFI [NP396/D^b^]) from the NIH Tetramer Core at Emory University. For gp33/D^b^ monomer, PE/Cy7-streptavidin was used. For gp276/D^b^ and NP396/D^b^, R-PE-streptavidin was used. Fluorophore-conjugated streptavidin was added to 20 μg of monomer, in 10 additions of 3.18 μg, each 10 min after the other (at room temperature). Tetramer was then stored at 4 °C before use.

### Software

Flow cytometry data was acquired in FACSDiva and analyzed in FlowJo v7 and v10. Statistical calculations and graphing were performed using GraphPad Prism v9. Figures were generated in Adobe Illustrator 2023.

## QUANTIFICATION AND STATISTICAL ANALYSIS

All statistical analyses were performed in Prism, as specified in the figure legends. Unpaired, two-tailed Student’s t tests were applied to determine the difference between two independent groups. Paired, two-tailed Student’s t tests were applied to determine the difference between two related groups (i.e. co-transferred populations within the same recipient animal). Where no p value is provided for a relevant comparison, the result was not significant. Infrequently, a ratio could not be calculated due to no cells in the *Il7ra*-deficient group remaining (i.e. 100 *Il7ra*-sufficient cells divided by 0 *Il7ra*-deficient treated cells). In this case, the ratio was set conservatively at the minimum possible value (i.e. 100:0 becomes 100:1, or a ratio of 100). Infrequently, too few events (less than five events) were captured for accurate quantitation of downstream parameters (i.e. Ki67, BrdU, CD127 gMFI). In such cases, these values were excluded for these parameters. In rare cases of sickness, animals were excluded. Error bars represent the SEM.

## SUPPLEMENTAL INFORMATION

**Figure S1, related to Figures 1 and 2.**
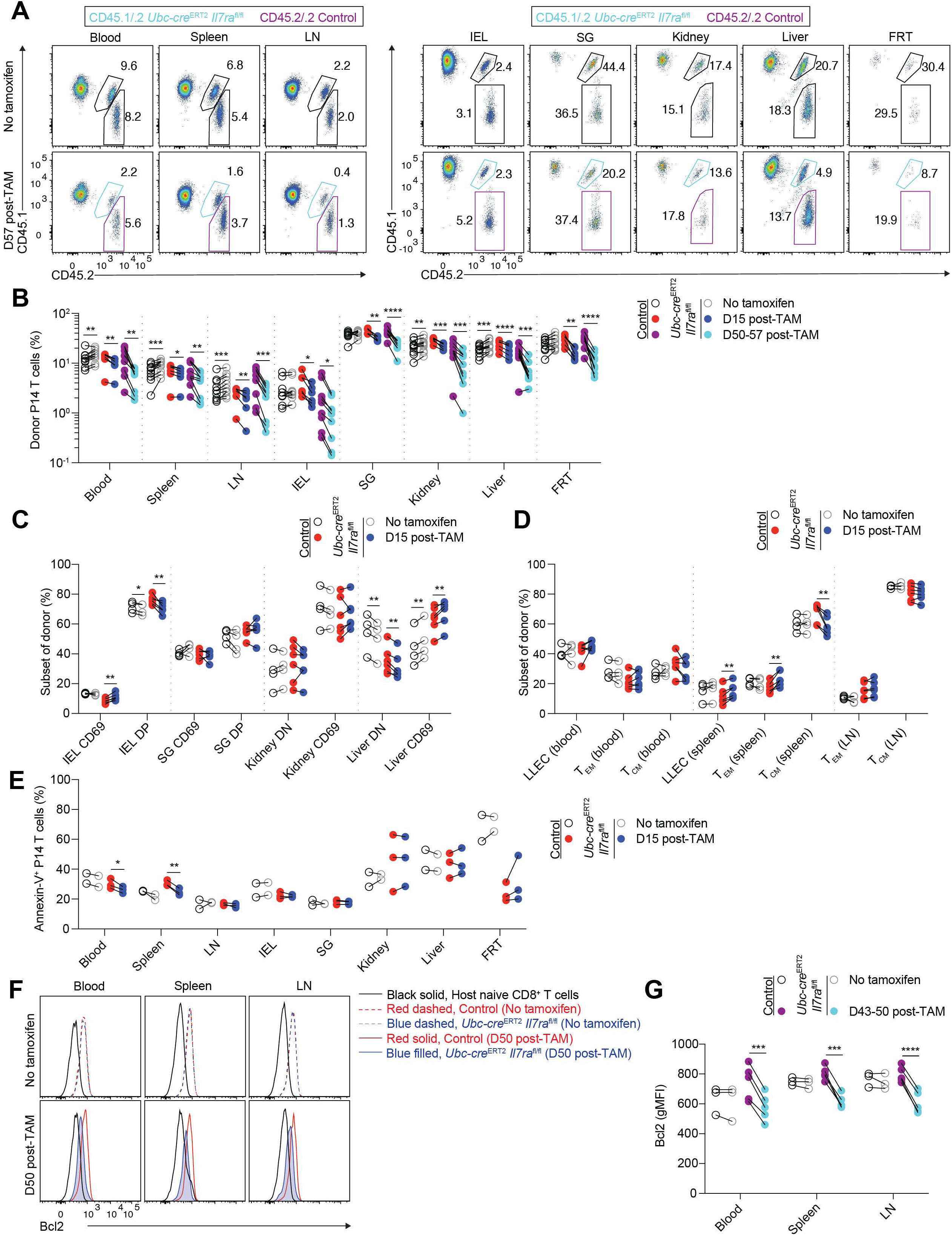
Loss of *Il7ra* from homeostatic memory cells reduces Bcl2 levels. As in Figures 1 and 2, congenic control (*Ubc-cre*^ERT2^ *Il7ra*^+/+^ or *Il7ra*^fl/fl^) and *Ubc-cre*^ERT2^ *Il7ra*^fl/fl^ P14 T cells were co-transferred to recipients, followed by LCMV-Armstrong infection one day later. After resting to memory phase (>28 days), some mice were given i.p. tamoxifen for 5 consecutive days to initiate Cre-mediated recombination of the floxed *Il7ra* allele. (A) Representative flow cytometry for congenic markers (CD45.2/.2 control and CD45.1/.2 *Ubc-cre*^ERT2^ *Il7ra*^fl/fl^) and quantitation of the (B) percent (of CD8^+^ T cells) as in (A), (C) proportions of NLT memory P14 subsets, (D) proportions of LLEC, T_EM_, and T_CM_, and (E) proportion of annexin V-staining control and *Ubc-cre*^ERT2^ *Il7ra*^fl/fl^ memory P14 T cells from untreated and tamoxifen-treated mice. (F) Representative flow cytometry for Bcl2 expression and (G) quantitation of Bcl2 gMFI on control and *Ubc-cre*^ERT2^ *Il7ra*^fl/fl^ memory P14 T cells from untreated and tamoxifen-treated mice as in (F). Data are (A,F) representative of 2-3 experiments (n=3-9/group), (B-D) compiled from 2-5 experiments (n=6-9/group), (E) from 1 experiment (n=2-3/group), or (G) compiled from 2 experiments (n=3-5/group). Error is expressed as ± S.E.M. Paired Student’s t tests. * p<0.05. ** p<0.01. *** p<0.001. **** p<0.0001.

**Figure S2, related to Figure 1.**
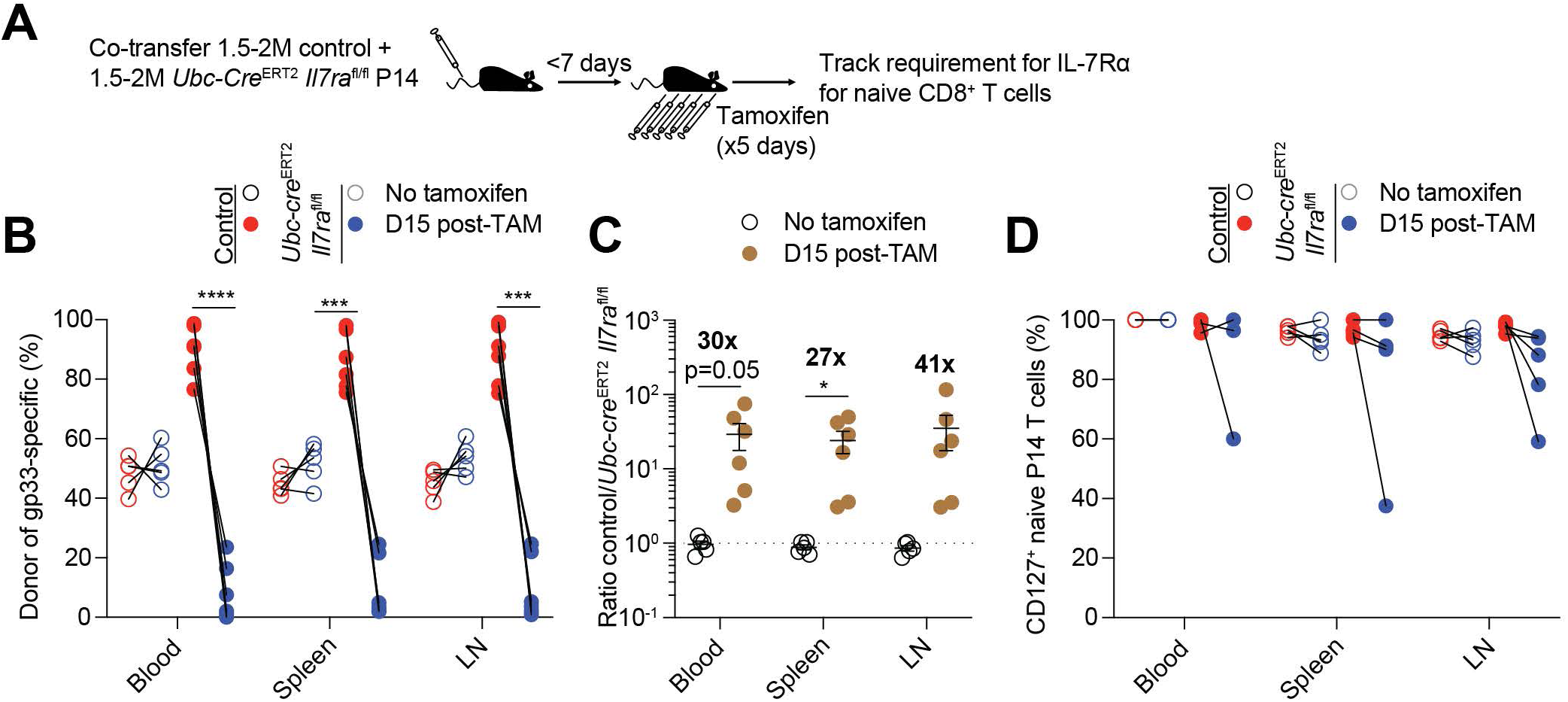
Naïve P14 T cells are profoundly dependent on IL-7Rα. (A) 1.5-2 million congenic control (*Ubc-cre*^ERT2^ *Il7ra*^+/+^ or *Il7ra*^fl/fl^) and *Ubc-cre*^ERT2^ *Il7ra*^fl/fl^ P14 T cells were co-transferred to naive recipients. Within 1 week of transfer, some mice were given i.p. tamoxifen for 5 consecutive days to initiate Cre-mediated recombination of the floxed *Il7ra* allele and tracked over time via gp33/D^b^ tetramer staining and congenic markers. (B-D) Quantitation of the (B) percent (of gp33/D^b^ tetramer-binding CD8^+^ T cells), (C) ratio, and (D) percent CD127^+^ of control and *Ubc-cre*^ERT2^ *Il7ra*^fl/fl^ naïve P14 T cells from untreated and tamoxifen-treated mice. Data are compiled from 3 experiments (n=3-6/group). Error is expressed as ± S.E.M. Unpaired (C) or paired (B,D) Student’s t tests. * p<0.05. ** p<0.01. *** p<0.001. **** p<0.0001.

**Figure S3, related to Figure 2.**
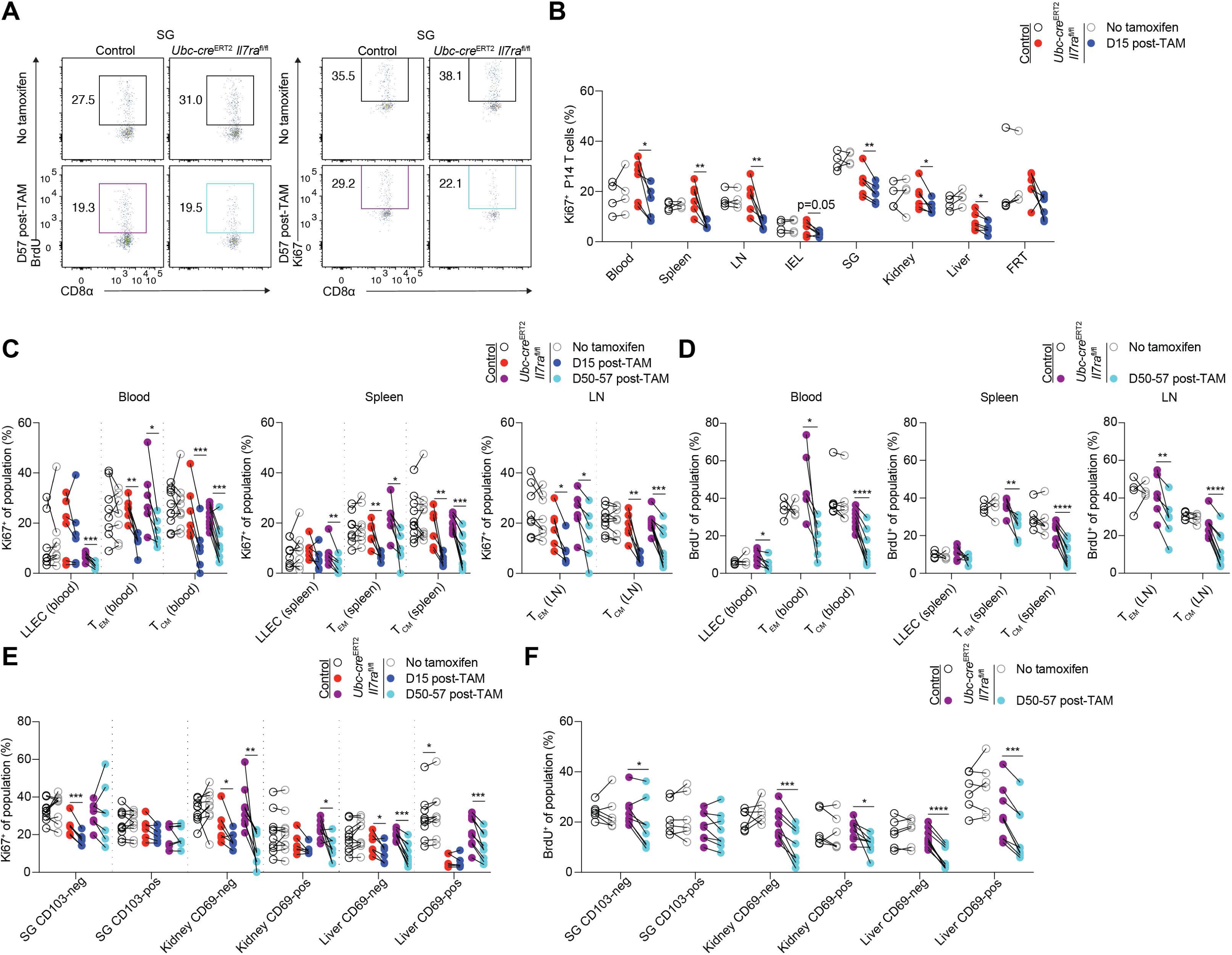
Loss of IL-7Rα affects circulating and resident memory P14 subsets. As in Figures 1 and 2, congenic control (*Ubc-cre*^ERT2^ *Il7ra*^+/+^ or *Il7ra*^fl/fl^) and *Ubc-cre*^ERT2^ *Il7ra*^fl/fl^ P14 T cells were co-transferred to recipients, followed by LCMV-Armstrong infection one day later. After resting to memory phase (>28 days), some mice were given i.p. tamoxifen for 5 consecutive days to initiate Cre-mediated recombination of the floxed *Il7ra* allele. (A) Representative flow cytometry for Ki67 expression and BrdU incorporation from the salivary gland and (B) quantitation of the proportion of Ki67-expressing control and *Ubc-cre*^ERT2^ *Il7ra*^fl/fl^ memory P14 T cells as in (A and Fig. 2A) from untreated and tamoxifen-treated mice. (C and D) Quantitation of the proportion of (C) Ki67-expressing and (D) BrdU-incorporating circulating memory P14 subsets from untreated and tamoxifen-treated mice. (E and F) Quantitation of the proportion of (E) Ki67- expressing and (F) BrdU-incorporating NLT memory P14 subsets from untreated and tamoxifen-treated mice. Data are compiled from 2-5 experiments (n=4-10/group). Error is expressed as ± S.E.M. Paired Student’s t tests. * p<0.05. ** p<0.01. *** p<0.001. **** p<0.0001.

**Figure S4, related to Figure 3.**
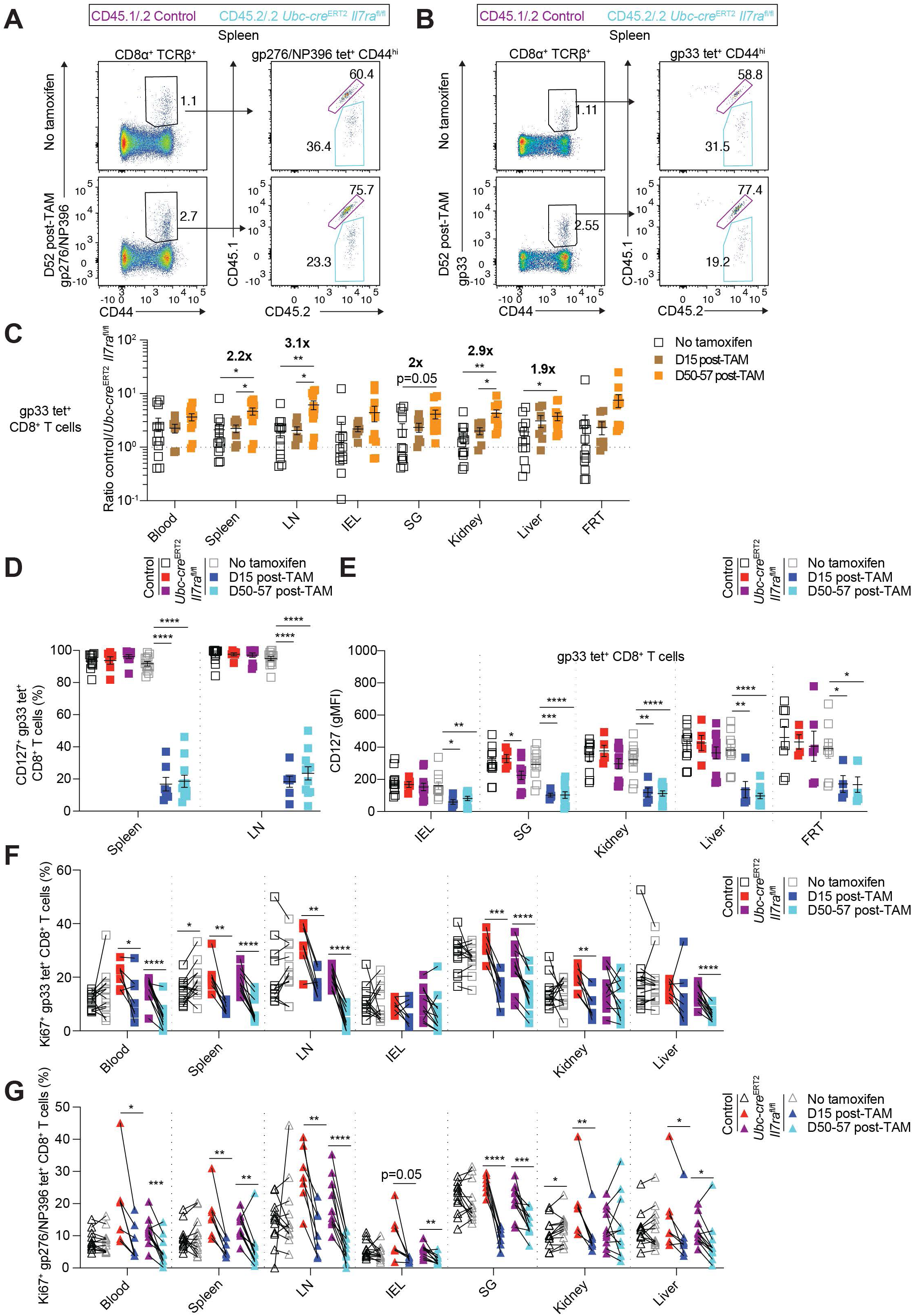
Antigen-specific memory CD8^+^ T cells are resilient to loss of *Il7ra* during memory homeostasis. As in Figure 3, congenic *Il7ra*^fl/fl^ *Ubc-cre*^ERT2^-negative (control) and -positive bone marrow was co-transferred to lethally irradiated recipients. After >8 weeks of reconstitution, animals were then infected with LCMV-Armstrong. After resting to memory phase (>28 days), some mice were given i.p. tamoxifen for 5 consecutive days to initiate Cre-mediated recombination of the floxed *Il7ra* allele and tracked over time via congenic markers. (A and B) Representative staining with (A) pooled gp276/D^b^ and NP396/D^b^ tetramers and (B) gp33/D^b^ tetramer for splenic memory CD8^+^ T cells and CD45.1/CD45.2 congenic gating (CD45.1/.2 *Il7ra*^fl/fl^ and CD45.2/.2 *Ubc-cre*^ERT2^ *Il7ra*^fl/fl^) from untreated and tamoxifen-treated bone marrow chimeras. (C) Ratio of control and *Ubc-cre*^ERT2^ *Il7ra*^fl/fl^ gp33/D^b^ tetramer-binding memory cells from untreated and tamoxifen-treated bone marrow chimeras gated as in (B). (D-F) Quantitation of the (D) proportion CD127^+^, (E) CD127 gMFI, and (F) proportion of Ki67-expressing cells for control and *Ubc-cre*^ERT2^ *Il7ra*^fl/fl^ gp33/D^b^ tetramer-binding memory cells from untreated and tamoxifen-treated bone marrow chimeras. (G) Quantitation of the proportion of Ki67-expressing cells for control and *Ubc-cre*^ERT2^ *Il7ra*^fl/fl^ gp276/D^b^/NP396/D^b^ tetramer-binding memory cells from untreated and tamoxifen-treated bone marrow chimeras (shares some data points with 3H). Data are (A,B) representative of 4 experiments (n=6-12/group) or (C-G) compiled from 3-7 experiments (n=6-12/group). Error is expressed as ± S.E.M. Unpaired (C-E) and paired Student’s t tests (F,G). * p<0.05. ** p<0.01. *** p<0.001. **** p<0.0001.

**Figure S5, related to Figure 5.**
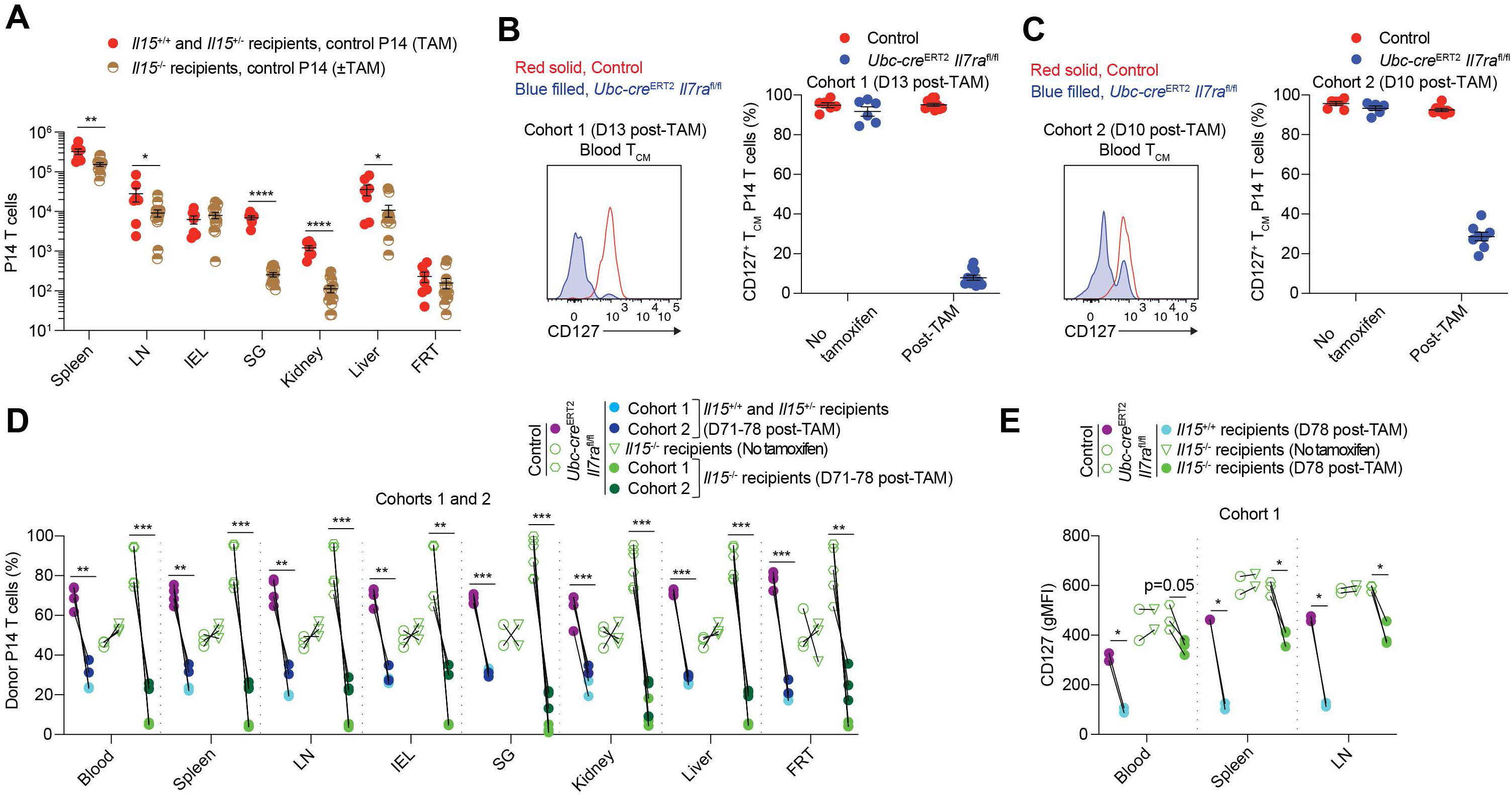
Redundancy between IL-7 and IL-15 affords resilience to loss of individual cytokines, part 1. (A, D, and E) As in Figure 5, congenic control (*Ubc-cre*^ERT2^ *Il7ra*^+/+^ or *Il7ra*^fl/fl^) and *Ubc-cre*^ERT2^ *Il7ra*^fl/fl^ P14 T cells were co-transferred to IL-15-sufficient (*Il15*^+/+^ and *Il15*^+/-^) or -deficient recipients, followed by LCMV-Armstrong infection one day later. After resting to memory phase (>28 days), some mice were given i.p. tamoxifen for 5 consecutive days to initiate Cre-mediated recombination of the floxed *Il7ra* allele and tracked over time in the blood via gp33/D^b^ tetramer staining and congenic markers. (A) Quantitation of control memory P14 T cells from untreated and tamoxifen-treated mice at D109-116 post-LCMV. (B and C) Congenic control and *Ubc-cre*^ERT2^ *Il7ra*^fl/fl^ P14 T cells were co-transferred to recipients (here wildtype, but also concurrently to IL-15- deficient recipients), followed by LCMV-Armstrong infection one day later. After resting to memory phase (>28 days), some mice were given i.p. tamoxifen for 5 consecutive days to initiate Cre-mediated recombination of the floxed *Il7ra* allele and tracked over time in the blood. Left, representative flow cytometry for CD127 expression and right, quantitation of the proportion of blood CD127^+^ T_CM_ P14 for (B) high efficiency Cohort 1 and (C) moderate efficiency Cohort 2. (D and E) As in Fig. S5A, Cohort 1 and 2 control and *Ubc-cre*^ERT2^ *Il7ra*^fl/fl^ memory P14 T cells were quantitated for (D) percent donor (of gp33/D^b^ tetramer-binding CD8^+^ T cells) and (E) CD127 gMFI from untreated and tamoxifen-treated mice. Data are compiled from (A) 3 experiments (n=6-13/group), (B,C) one experiment representative of 3-4 experiments (n=6-10/group), (D) 2 experiments (n=2-6/group), or (E) 1 experiment (n=2-3/group). Error is expressed as ± S.E.M. Unpaired (A) and paired Student’s t tests (D, E). * p<0.05. ** p<0.01. *** p<0.001. **** p<0.0001.

**Figure S6, related to Figure 5.**
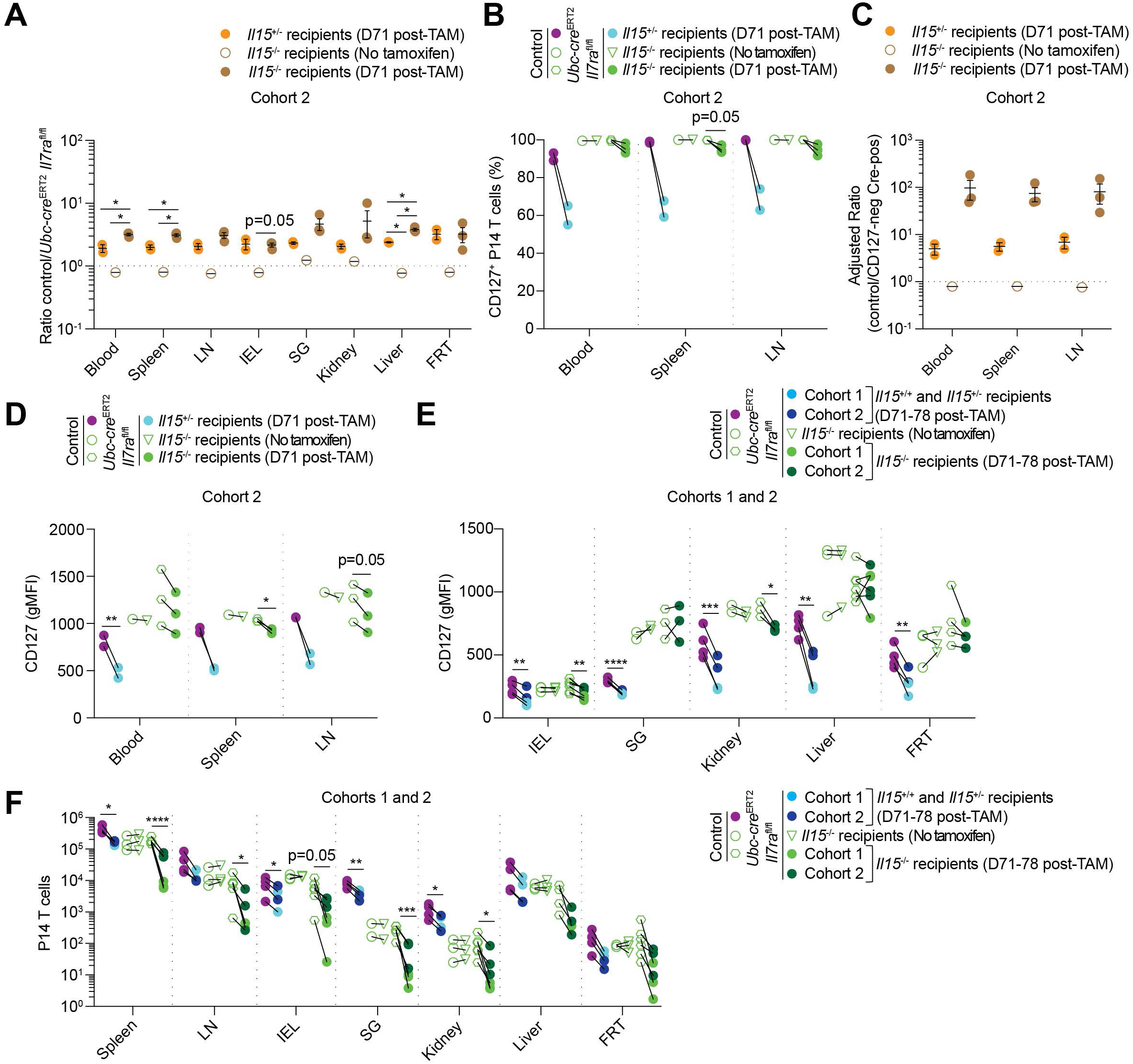
Redundancy between IL-7 and IL-15 affords resilience to loss of individual cytokines, part 2. As in Figures 5 and S5, (A-D) Cohort 2 control and *Ubc-cre*^ERT2^ *Il7ra*^fl/fl^ memory P14 T cells were quantitated for (A) ratio, (B) proportion CD127^+^, (C) adjusted ratio of control to CD127-negative *Ubc-cre*^ERT2^ *Il7ra*^fl/fl^ memory P14 T cells, and (D) CD127 gMFI from untreated and tamoxifen-treated mice. (E) Cohort 1 and 2 control and *Ubc-cre*^ERT2^ *Il7ra*^fl/fl^ memory P14 T cells were quantitated for CD127 gMFI from untreated and tamoxifen-treated mice. (F) Quantitation of Cohort 1 and 2 control and *Ubc-cre*^ERT2^ *Il7ra*^fl/fl^ memory P14 T cells from untreated and tamoxifen-treated mice (some data points are also shown in Fig. S5A). Data are (A-D) from 1 experiment (n=1-3/group) or (E,F) compiled from 1-2 experiments (n=2-6/group). Error is expressed as ± S.E.M. Unpaired (A,C) and paired Student’s t tests (B,D-F). * p<0.05. ** p<0.01. *** p<0.001. **** p<0.0001.

**Figure S7, related to Figure 6.**
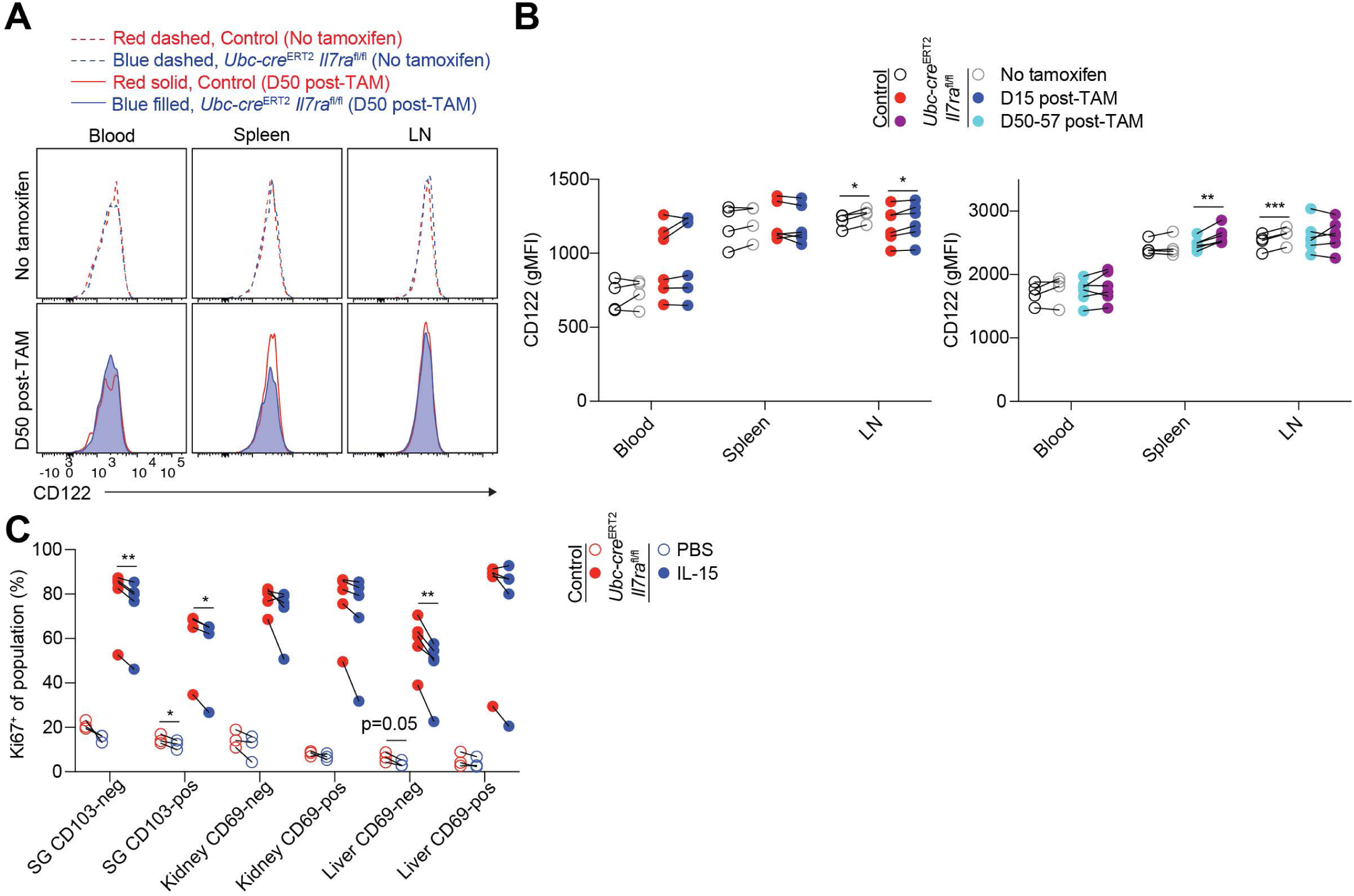
IL-7Rα regulates responsiveness to IL-15. (A and B) As in Figures 1 and 2, congenic control (*Ubc-cre*^ERT2^ *Il7ra*^+/+^ or *Il7ra*^fl/fl^) and *Ubc-cre*^ERT2^ *Il7ra*^fl/fl^ P14 T cells were co-transferred to recipients, followed by LCMV-Armstrong infection one day later. After resting to memory phase (>28 days), some mice were given i.p. tamoxifen for 5 consecutive days to initiate Cre-mediated recombination of the floxed *Il7ra* allele. (A) Representative flow cytometry for CD122 expression and (B) quantitation of CD122 gMFI as in (A) for control and *Ubc-cre*^ERT2^ *Il7ra*^fl/fl^ P14 T cells from untreated and tamoxifen-treated recipients. Left, D15 post-TAM timepoint. Right, D50-57 post-TAM timepoint. (C) As in Figure 6G-H, quantitation of the proportion of Ki67-expressing NLT memory subset P14 T cells from PBS- and IL-15-treated mice (all previously treated with tamoxifen). Data are (A) representative of 2 experiments (n=4-6/group) or (B,C) compiled from 2-3 experiments (n=4-12/group). Error is expressed as ± S.E.M. Paired Student’s t tests. * p<0.05. ** p<0.01. *** p<0.001. **** p<0.0001.

## REFERENCES

1. Masopust, D., Vezys, V., Marzo, A.L., and Lefrancois, L. (2001). Preferential localization of effector memory cells in nonlymphoid tissue. Science 291, 2413–2417. 10.1126/science.1058867

2. Jameson, S.C. and Masopust, D. (2018). Understanding Subset Diversity in T Cell Memory. Immunity 48, 214–226. 10.1016/j.immuni.2018.02.010

3. Masopust, D. and Soerens, A.G. (2019). Tissue-Resident T Cells and Other Resident Leukocytes. Ann. Rev. Immunol. 37, 521–546. 10.1146/annurev-immunol-042617-053214

4. Steinert, E.M., Schenkel, J.M., Fraser, K.A., Beura, L.K., Manlove, L.S., Igyártó, B.Z., Southern, P.J., Masopust, D. (2015). Quantifying Memory CD8 T Cells Reveals Regionalization of Immunosurveillance. Cell 161, 737–748. 10.1016/j.cell.2015.03.031

5. Wijeyesinghe, S., Beura, L.K., Pierson, M.J., Stolley, J.M., Adam, O.A., Ruscher, R., Steinert, E.M., Rosato, P.C., Vezys, V., Masopust, D. (2021). Expansible residence decentralizes immune homeostasis. Nature 592, 457–462. 10.1038/s41586-021-03351-3

6. Mackay, L.K., Rahimpour, A., Ma, J.Z., Collins, N., Stock, A.T., Hafon, M.-L., Vega-Ramos, J., Lauzurica, P., Mueller, S.N., Stefanovic, T., et al. (2013). The developmental pathway for CD103(+)CD8+ tissue-resident memory T cells of skin. Nat. Immunol. 14, 1294–1301. 10.1038/ni.2744

7. Milner, J.J., Toma, C., Yu, B., Zhang, K., Omilusik, K., Phan, A.T., Wang, D., Getzler, A. J., Nguyen, T., Crotty, S., et al. (2017). Runx3 programs CD8 + T cell residency in non-lymphoid tissues and tumours. Nature 552, 253–257. 10.1038/nature24993

8. Kumar, B.V., Ma, W., Miron, M., Granot, T., Guyer, R.S., Carpenter, D.J., Senda, T., Sun, X., Ho, S.-H., Lerner, H., et al. (2017). Human Tissue-Resident Memory T Cells Are Defined by Core Transcriptional and Functional Signatures in Lymphoid and Mucosal Sites. Cell Rep. 20, 2921–2934. 10.1016/j.celrep.2017.08.078

9. Frizzell, H., Fonseca, R., Christo, S.N., Evrard, M., Cruz-Gomez, S., Zanluqui, N.G., von Scheidt, B., Freestone, D., Park, S.L., McWilliam, H.E.G., et al. (2020). Organ-specific isoform selection of fatty acid-binding proteins in tissue-resident lymphocytes. Sci. Immunol. 5, eaay9283. 10.1126/sciimmunol.aay9283

10. Christo, S.N., Evrard, M., Park, S.L., Gandolfo, L.C., Burn, T.N., Fonseca, R., Newman, D.M., Alexandre, Y.O., Collins, N., Zamudio, N.M., et al. (2021). Discrete tissue microenvironments instruct diversity in resident memory T cell function and plasticity. Nat. Immunol. 22, 1140–1151. 10.1038/s41590-021-01004-1

11. Crowl, J.T., Heeg, M., Ferry, A., Milner, J.J., Omilusik, K.D., Toma, C., He, Z., Chang, J.T., and Goldrath, A.W. (2022). Tissue-resident memory CD8 + T cells possess unique transcriptional, epigenetic and functional adaptations to different tissue environments. Nat. Immunol. 23, 1121–1131. 10.1038/s41590-022-01229-8

12. Poon, M.M.L., Caron, D.P., Wang, Z., Wells, S.B., Chen, D., Meng, W., Szabo, P.A., Lam, N., Kubota, M., Matsumoto, R., et al. (2023). Tissue adaptation and clonal segregation of human memory T cells in barrier sites. Nat. Immunol. 24, 309–319. 10.1038/s41590-022-01395-9

13. Gebhardt, T., Wakim, L.M., Eidsmo, L., Reading, P.C., Heath, W.R., and Carbone, F.R. (2009). Memory T cells in nonlymphoid tissue that provide enhanced local immunity during infection with herpes simplex virus. Nat. Immunol. 10, 524–530. 10.1038/ni.1718

14. Jiang, X., Clark, R.A., Liu, L., Wagers, A.J., Fuhlbrigge, R.C., and Kupper, T.S. (2012). Skin infection generates non-migratory memory CD8+ T(RM) cells providing global skin immunity. Nature 483, 227–231. 10.1038/nature10851

15. Beura, L.K., Mitchell, J.S., Thompson, E.A., Schenkel, J.M., Mohammed, J., Wijeyesinghe, S., Fonseca, R., Burbach, B.J., Hickman, H.D., Vezys, V., et al. (2018). Intravital mucosal imaging of CD8 + resident memory T cells shows tissue-autonomous recall responses that amplify secondary memory. Nat. Immunol. 19, 173–182. 10.1038/s41590-017-0029-3

16. Park, S.L., Zaid, A., Hor, J.L., Christo, S.N., Prier, J.E., Davies, B., Alexandre, Y.O., Gregory, J.L., Russell, T.A., Gebhardt, T., et al. (2018). Local proliferation maintains a stable pool of tissue-resident memory T cells after antiviral recall responses. Nat. Immunol. 19, 183–191. 10.1038/s41590-017-0027-5

17. Park, S.L., Buzzai, A., Rautela, J., Hor, J.L., Hochheiser, K., Effern, M., McBain, N., Wagner, T., Edwards, J., McConville, R., et al. (2019). Tissue-resident memory CD8 + T cells promote melanoma-immune equilibrium in skin. Nature 565, 366–371. 10.1038/s41586-019-0958-0

18. Lodolce, J.P., Boone, D.L., Chai, S., Swain, R.E., Dassopoulos, T., Trettin, S., and Ma, A. (1998). IL-15 receptor maintains lymphoid homeostasis by supporting lymphocyte homing and proliferation. Immunity 9, 669–676. 10.1016/s1074-7613(00)80664-0

19. Kennedy, M.K., Glaccum, M., Brown, S.N., Butz, E.A., Viney, J.L., Embers, M., Matsuki, N., Charrier, K., Sedger, L., Willis, C.R., et al. (2000). Reversible defects in natural killer and memory CD8 T cell lineages in interleukin 15-deficient mice. J. Exp. Med. 191, 771–780. 10.1084/jem.191.5.771

20. Schluns, K.S., Kieper, W.C., Jameson, S.C., and Lefrancois, L. (2000). Interleukin-7 mediates the homeostasis of naïve and memory CD8 T cells in vivo. Nat. Immunol. 1, 426–432. 10.1038/80868

21. Becker, T.C., Wherry, E.J., Boone, D., Murali-Krishna, K., Antia, R., Ma, A., and Ahmed, R. (2002). Interleukin 15 is required for proliferative renewal of virus-specific memory CD8 T cells. J. Exp. Med. 195, 1541–1548. 10.1084/jem.20020369

22. Goldrath, A.W., Sivakumar, P.V., Glaccum, M., Kennedy, M.K., Bevan, M.J., Benoist, C., Mathis, D., and Butz, E.A. (2002). Cytokine requirements for acute and Basal homeostatic proliferation of naive and memory CD8+ T cells. J. Exp. Med. 195, 1515–1522. 10.1084/jem.20020033

23. Judge, A.D., Zhang, X., Fujii, H., Surh, C.D., and Sprent, J. (2002). Interleukin 15 controls both proliferation and survival of a subset of memory-phenotype CD8(+) T cells. J. Exp. Med. 196, 935–946. 10.1084/jem.20020772

24. Schluns, K.S., Williams, K., Ma, A., Zheng, X.X., and Lefrançois, L. (2002). Cutting edge: requirement for IL-15 in the generation of primary and memory antigen-specific CD8 T cells. J. Immunol. 168, 4827–4831. 10.4049/jimmunol.168.10.4827

25. Sandau, M.M., Kohlmeier, J.E., Woodland, D.L., and Jameson, S.C. (2010). IL-15 regulates both quantitative and qualitative features of the memory CD8 T cell pool. J. Immunol. 184, 35–44. 10.4049/jimmunol.0803355

26. Masopust, D., Vezys, V., Wherry, E.J., Barber, D.L., and Ahmed, R. (2006). Cutting edge: gut microenvironment promotes differentiation of a unique memory CD8 T cell population. J. Immunol. 176, 2079–2083. 10.4049/jimmunol.176.4.2079

27. Jarjour, N.N., Wanhainen, K.M., Peng, C., Gavil, N.V., Maurice, N.J., Borges da Silva, H., Martinez, R.J., Dalzell, T.S., Huggins, M.A., Masopust, D., et al. (2022). Responsiveness to interleukin-15 therapy is shared between tissue-resident and circulating memory CD8 + T cell subsets. Proc. Natl. Acad. Sci. U.S.A. 119, e2209021119. 10.1073/pnas.2209021119

28. Schenkel, J.M., Fraser, K.A., Casey, K.A., Beura, L.K., Pauken, K.E., Vezys, V., and Masopust, D. (2016). IL-15-Independent Maintenance of Tissue-Resident and Boosted Effector Memory CD8 T Cells. J. Immunol. 196, 3920–3926. 10.4049/jimmunol.1502337

29. Verbist, K.C., Field, M.B., and Klonowski, K.D. (2011). Cutting edge: IL-15-independent maintenance of mucosally generated memory CD8 T cells. J. Immunol. 186, 6667–6671. 10.4049/jimmunol.1004022

30. Kaech, S.M. and Cui, W. (2012). Transcriptional control of effector and memory CD8+ T cell differentiation. Nat. Rev. Immunol. 12, 749–761. 10.1038/nri3307

31. Carrette, F. and Surh, C.D. (2012). IL-7 signaling and CD127 receptor regulation in the control of T cell homeostasis. Semin. Immunol. 24, 209–217. 10.1016/j.smim.2012.04.010

32. Hashimoto, M., Im, S.J., Araki, K., and Ahmed, R. (2019). Cytokine-Mediated Regulation of CD8 T-Cell Responses During Acute and Chronic Viral Infection. Cold Spring Harb. Perspect. Biol. 11, a028464. 10.1101/cshperspect.a028464

33. Kawabe, T., Yi, J., and Sprent, J. (2021). Homeostasis of Naive and Memory T Lymphocytes. Cold Spring Harb. Perspect. Biol. 13, a037879. 10.1101/cshperspect.a037879

34. Adachi, T., Kobayashi, T., Sugihara, E., Yamada, T., Ikuta, K., Pittaluga, S., Saya, H., Amagai, M., and Nagao, K. (2015). Hair follicle-derived IL-7 and IL-15 mediate skin-resident memory T cell homeostasis and lymphoma. Nat. Med. 21, 1272–1279. 10.1038/nm.3962

35. Buentke, E., Mathiot, A., Tolaini, M., Di Santo, J., Zamoyska, R., and Seddon, B. (2006). Do CD8 effector cells need IL-7R expression to become resting memory cells? Blood 108, 1949–1956. 10.1182/blood-2006-04-016857

36. Kaech, S.M., Tan, J.T., Wherry, E.J., Konieczny, B.T., Surh, C.D., and Ahmed, R. (2003). Selective expression of the interleukin 7 receptor identifies effector CD8 T cells that give rise to long-lived memory cells. Nat. Immunol. 4, 1191–1198. 10.1038/ni1009

37. Lenz, D.C., Kurz, S.K., Lemmens, E., Schoenberger, S.P., Sprent, J., Oldstone, M.B.A., and Homann, D. (2004). IL-7 regulates basal homeostatic proliferation of antiviral CD4+ T cell memory. Proc. Natl. Acad. Sci. U.S.A. 101, 9357–9362. 10.1073/pnas.0400640101

38. Klonowski, K.D., Williams, K.J., Marzo, A.L., and Lefrançois, L. (2006). Cutting edge: IL-7- independent regulation of IL-7 receptor alpha expression and memory CD8 T cell development. J. Immunol. 177, 4247–4251. 10.4049/jimmunol.177.7.4247

39. Carrio, R., Rolle, C.E., and Malek, T.R. (2007). Non-redundant role for IL-7R signaling for the survival of CD8+ memory T cells. Eur. J. Immunol. 37, 3078–3088. 10.1002/eji.200737585

40. Osborne, L.C., Dhanji, S., Snow, J.W., Priatel, J.J., Ma, M.C., Miners, M.J., Teh, H.-S., Goldsmith, M.A., and Abraham, N. (2007). Impaired CD8 T cell memory and CD4 T cell primary responses in IL-7R alpha mutant mice. J. Exp. Med. 204, 619–631. 10.1084/jem.20061871

41. Rubinstein, M.P., Lind, N.A., Purton, J.F., Filippou, P., Best, J.A., McGhee, P.A., Surh, C.D., and Goldrath, A.W. (2008). IL-7 and IL-15 differentially regulate CD8+ T-cell subsets during contraction of the immune response. Blood 112, 3704–3712. 10.1182/blood-2008-06-160945

42. Park, J.-H., Yu, Q., Erman, B., Appelbaum, J.S., Montoya-Durango, D., Grimes, H.L., and Singer, A. (2004). Suppression of IL7Ralpha transcription by IL-7 and other prosurvival cytokines: a novel mechanism for maximizing IL-7-dependent T cell survival. Immunity 21, 289–302. 10.1016/j.immuni.2004.07.016

43. Surh, C.D. and Sprent, J. (2005). Regulation of mature T cell homeostasis. Semin. Immunol. 17, 183–191. 10.1016/j.smim.2005.02.007

44. Jacobs, S.R., Michalek, R.D., and Rathmell, J.C. (2010). IL-7 Is Essential for Homeostatic Control of T Cell Metabolism In Vivo. J. Immunol. 184, 3461–3469. 10.4049/jimmunol.0902593

45. Anderson, K.G., Mayer-Barber, K., Sung, H., Beura, L., James, B.R., Taylor, J.J., Qunaj, L., Griffith, T.S., Vezys, V., Barber, D.L., et al. (2014). Intravascular staining for discrimination of vascular and tissue leukocytes. Nat. Protoc. 9, 209–222. 10.1038/nprot.2014.005

46. Grayson, J.M., Zajac, A.J., Altman, J.D., and Ahmed, R. (2000). Cutting edge: increased expression of Bcl-2 in antigen-specific memory CD8+ T cells. J. Immunol. 164, 3950–3954. 10.4049/jimmunol.164.8.3950

47. Williams, M.A., Tyznik, A.J., and Bevan, M.J. (2006). Interleukin-2 signals during priming are required for secondary expansion of CD8+ memory T cells. Nature 441, 890–893. 10.1038/nature04790

48. Kaech, S.M. and Ahmed, R. (2001). Memory CD8+ T cell differentiation: initial antigen encounter triggers a developmental program in naïve cells. Nat. Immunol. 2, 415–422. 10.1038/87720

49. Tan, J.T., Ernst, B., Kieper, W.C., LeRoy, E., Sprent, J., and Surh, C.D. (2002). Interleukin (IL)-15 and IL-7 jointly regulate homeostatic proliferation of memory phenotype CD8+ cells but are not required for memory phenotype CD4+ cells. J. Exp. Med. 195, 1523–1532. 10.1084/jem.20020066

50. Rubinstein, M.P., Kovar, M., Purton, J.F., Cho, J.-H., Boyman, O., Surh, C.D., and Sprent, J. (2006). Converting IL-15 to a superagonist by binding to soluble IL-15R{alpha}. Proc. Natl. Acad. Sci. U.S.A. 103, 9166–9171. 10.1073/pnas.0600240103

51. Stoklasek, T.A., Schluns, K.S., and Lefrançois, L. (2006). Combined IL-15/IL-15Ralpha immunotherapy maximizes IL-15 activity in vivo. J. Immunol. 177, 6072–6080. 10.4049/jimmunol.177.9.6072

52. Boyman, O., Ramsey, C., Kim, D.M., Sprent, J., and Surh, C.D. (2008). IL-7/anti-IL-7 mAb complexes restore T cell development and induce homeostatic T Cell expansion without lymphopenia. J. Immunol. 180, 7265–7275. 10.4049/jimmunol.180.11.7265

53. Zhang, X., Sun, S., Hwang, I., Tough, D., and Sprent, J. (1998). Potent and selective stimulation of memory-phenotype CD8+ T cells in vivo by IL-15. Immunity 8, 591–599. 10.1016/s1074-7613(00)80564-6

54. Link, A., Vogt, T.K., Favre, S., Britschgi, M.R., Acha-Orbea, H., Hinz, B., Cyster, J.G., and Luther, S.A. (2007). Fibroblastic reticular cells in lymph nodes regulate the homeostasis of naive T cells. Nat. Immunol. 8, 1255–1265. 10.1038/ni1513

55. Onder, L., Narang, P., Scandella, E., Chai, Q., Iolyeva, M., Hoorweg, K., Halin, C., Richie, E., Kaye, P., Westermann, J., et al. (2012). IL-7-producing stromal cells are critical for lymph node remodeling. Blood 120, 4675–4683. 10.1182/blood-2012-03-416859

56. Sheikh, A., Jackson, J., Shim, H. B., Yau, C., Seo, J. H., and Abraham, N. (2022). Selective dependence on IL-7 for antigen-specific CD8 T cell responses during airway influenza infection. Sci. Rep. 12, 135. 10.1038/s41598-021-03936-y

57. Ikuta, K., Hara, T., Abe, S., Asahi, T., Takami, D., and Cui, G. (2021). The Roles of IL-7 and IL-15 in Niches for Lymphocyte Progenitors and Immune Cells in Lymphoid Organs. Curr. Top. Microbiol. Immunol. 434, 83–101. 10.1007/978-3-030-86016-5_4

58. McCaughtry, T.M., Etzensperger, R., Alag, A., Tai, X., Kurtulus, S., Park, J.-H., Grinberg, A., Love, P., Feigenbaum, L., Erman, B., et al. (2012). Conditional deletion of cytokine receptor chains reveals that IL-7 and IL-15 specify CD8 cytotoxic lineage fate in the thymus. J. Exp. Med. 209, 2263–2276. 10.1084/jem.20121505

59. Pircher, H., Mak, T.W., Lang, R., Ballhausen, W., Rüedi, E., Hengartner, H., Zinkernagel, R. M., and Bürki, K. (1987). T cell tolerance to Mlsa encoded antigens in T cell receptor V beta 8.1 chain transgenic mice. EMBO J. 8, 719–727. 10.1002/j.1460-2075.1989.tb03431.x

60. Heffner, C.S., Herbert, P., Babiuk, R.P., Sharma, Y., Rockwood, S.F., Donahue, L.R., Eppig, J.T., and Murray, S.A. (2012). Supporting conditional mouse mutagenesis with a comprehensive cre characterization resource. Nat. Comm. 3, 1218. 10.1038/ncomms2186

61. Burrack, K.S., Huggins, M.A., Taras, E., Dougherty, P., Henzler, C.M., Yang, R., Alter, S., Jeng, E.K., Wong, H.C., Felices, M., et al. (2018). Interleukin-15 Complex Treatment Protects Mice from Cerebral Malaria by Inducing Interleukin-10-Producing Natural Killer Cells. Immunity 48, 760–772. 10.1016/j.immuni.2018.03.012

